# Pervasive translation of intergenic open reading frames and *de novo* gene emergence in *Arabidopsis*

**DOI:** 10.1101/2025.09.04.674254

**Authors:** Claire Patiou, Christelle Blassiau, Isabelle Hatin, Lars A. Eicholt, Enora Corler, Chloé Ponitzki, Laurence Debacker, Erich Bornberg-Bauer, Olivier Namy, Sylvain Legrand, Vincent Castric, Eléonore Durand

**Author notes:** Author for Correspondence: Eléonore Durand, University of Lille, Lille, France.

## Abstract

Ancestrally non-genic sequences are now widely recognized as potential reservoirs for the *de novo* emergence of new genes. Across clades, some *de novo* genes were proven to have substantial phenotypic effects, and to contribute to the emergence of novel biological functions. Yet, little is still known about the starting material from which *de novo* genes emerge, especially in plants. To fill this gap, we generated Ribosome Profiling data from *Arabidopsis lyrata* and characterized the evolution of translated regions genome-wide across the Arabidopsis genus. Synteny analysis revealed 163 actively translated regions in *A. lyrata* whose coding potential (Open Reading Frames, ORFs or Coding DNA sequences, CDSs) has emerged *de novo* within the Arabidopsis genus. Most of these *de novo* translated regions were species- and even accession-specific, indicating their transient nature, with patterns of polymorphism consistent with neutral evolution in natural populations. They were also significantly shorter and less expressed than conserved protein-coding genes, and their GC content increased with phylogenetic conservation. Twenty-one of them belonged to previously annotated CDSs, and are therefore promising putative *de novo* genes, while most were located in intergenic regions and are thus newly discovered. Our results demonstrate the abundance of translation events outside of conserved CDSs, and their role as starting material for the emergence of novel genes in plants.

**Significance statement:** A central open question in genome evolution is how novel protein-coding genes arise from noncoding nucleotide DNA sequences and eventually contribute to the stable repertoires of “canonical” genes. Here, we focused specifically on the early stages of this important evolutionary process, whereby previously non-coding intergenic nucleotide sequences eventually acquire open reading frames carrying signatures of active translation. This phenomenon has been crucially under-studied so far, especially in plants. By combining ribosome profiling data and a careful genome comparison strategy among closely related Arabidopsis species, our results demonstrate the pervasive translation and *de novo* origin of a large number of small intergenic ORFs and illustrate their role as starting material for the emergence of novel genes in plants.

Key-words: de novo genes, Arabidopsis, pervasive translation, intergenic ORFs, ribosome profiling

## Introduction

According to François Jacob, “The probability of a functional protein appearing *de novo* by random association of amino acids is almost zero” (Jacob 1977). Yet, this historical paradigm was challenged by the gradual discovery, in multiple organisms, of genes that lacked homology to any other known protein-coding sequences, some of which were later characterized as having appeared *de novo* from ancestrally non-coding sequences (Begun et al. 2006; Levine et al. 2006; Cai et al. 2008; Knowles and McLysaght 2009; Xiao et al. 2009; Tautz and Domazet-Lošo 2011). The first *de novo* genes were proposed in *Drosophila* (Begun et al. 2006; Levine et al. 2006), yeast (Cai et al. 2008) and primates (Toll-Riera et al. 2008; Knowles and McLysaght 2009). Since these first discoveries, important *de novo* genes have been identified and functionally characterized in several taxa over the past decade, ranging from the anti-freeze *afgp* gene in cod (Baalsrud et al. 2018), to the DNA repair *BSC4* gene in yeast (Cai et al. 2008), the spermatogenesis related *goddard* gene in flies (Lange et al. 2021) or the oncopromoting factor *NCYM* gene in human cancer (Suenaga et al. 2014). In plants, the first identified *de novo* gene was the qua-quine-starch (*QQS*) gene in *A. thaliana*, which encodes a 59 amino-acid polypeptide involved in the regulation of starch synthesis in response to genetic and environmental perturbations (Li et al. 2009). A more recent example is the Sun Wu-Kong (*SWK*) *de novo* gene, which encodes a 65 amino-acid protein involved in seed germination under osmotic stress in *A. thaliana* (Jin et al. 2025). In rice, the *de novo* gene *GSE9* contributes to the shape of rice grains and thus the differences between *indica* and *japonica* varieties (Chen et al. 2023). Overall, *de novo* genes seem to be fairly common, with *e.g.* 782 putative *de novo* genes reported in *A. thaliana* (Li et al. 2016) and 175 in *O. sativa* (Zhang et al. 2019), although generalization is limited by the very few comprehensive genome-wide repertoires characterized in plants so far (Silveira et al. 2013; Cui et al. 2015; Li et al. 2016; Zhang et al. 2019; Song et al. 2022; Poretti et al. 2023; Cao et al. 2024; Li et al. 2025).

Our understanding of the early evolutionary processes by which *de novo* genes emerge from non-genic regions is limited by the focus of most studies on protein-coding sequences that had already been annotated as “genes”, and hence already harbor well-established “gene-like” characteristics (Ruiz-Orera et al. 2015; Vakirlis et al. 2018; Bornberg-Bauer et al. 2021; Peng and Zhao 2024). However, properties of *de novo* genes, particularly young *de novo* genes, appear to differ markedly from those of more established, evolutionarily conserved (“canonical”) genes. In particular, young *de nov*o genes often exhibit a simpler gene structure (fewer or no introns), lower expression and a narrower tissue specificity of expression as compared to conserved genes (Zhang et al. 2019, Jin et al. 2025). Little is known about the very first steps of *de novo* gene emergence, especially the properties of the ancestral non-coding sequences and how they eventually acquire their “gene-like” characteristics. So far, studies in yeast (Carvunis et al. 2012, Durand et al. 2019), and in *Mus musculus* (Ruiz-Orera et al. 2018, Schmitz et al. 2018) indicate that small, translated ORFs located in intergenic regions and without protein-coding gene orthologs in closely-related species could provide the basis for *de novo* gene emergence. Hundreds to thousands of such ORFs are found in yeast, flies and mice genomes, with a high rate of evolutionary turnover and mostly neutral patterns of molecular evolution (Carvunis et al. 2012, Durand et al. 2019, Ruiz-Orera et al. 2018, Grandchamp et al. 2023, Grandchamps et al. 2024). Hence, these ORFs can be viewed as a “reservoir” from which a few novel functional protein-coding genes might eventually emerge (Carvunis et al. 2012, Ruiz-Orera et al. 2018; Durand et al. 2019; Ruiz-Orera and Albà 2019).

The detailed annotation of translated intergenic regions that have emerged *de novo* is challenging for at least two reasons. First, evidence of translation is itself hard to obtain, precisely because *de novo* genes are not expected to harbor the hallmarks of canonical protein-coding sequences (such as greater length and sustained expression), and because their expression tends to be low (Lei et al. 2024). Obtaining translational evidence is nonetheless possible by using Ribosome Profiling, a powerful technology whereby RNA molecules bound to ribosomes are isolated and specifically sequenced, providing a detailed snapshot of translational activity at the genome-wide scale (Ingolia et al. 2009; Ingolia et al. 2014; Kiniry et al. 2020). In plants, only few high-quality ribosome profiling datasets are available, all from model species such as *O. sativa* (Zhu et al. 2023), *Zea mays* (Chotewutmontri et al. 2018) and *A. thaliana* (Hsu et al. 2016; Wu and Hsu 2022). Second, confidently asserting that a given coding sequence has emerged *de novo* requires the identification of a clear orthologous non-coding region in a sister species. However, homology detection lacks power beyond a certain degree of divergence, resulting in a potentially serious “phylostratigraphic bias” (Moyers and Zhang 2015; Moyers and Zhang 2016; Moyers and Zhang 2017; Moyers and Zhang 2018), whose importance has been debated (Domazet-Lošo et al. 2017; Weisman et al. 2020). A solid way to circumvent this problem, and increase the reliability of *de novo* gene detection, is to identify orthologous intergenic regions that have remained syntenic (Vakirlis et al. 2020). Because genomes can become drastically rearranged over the course of evolution, analysing synteny is typically challenging, if at all possible, among distantly related species, and instead performs best when comparing species that are closely related (Tang et al. 2008). Previous studies have tackled one of those two problems, either by using ribosome profiling sequencing to produce ribosome-protected mRNA fragments (RPFs) in a few distantly related species (Ruiz-Orera et al. 2018; Xiao et al. 2018), or by establishing detailed synteny relationships between closely related species, but without corresponding ribosome profiling data (Vakirlis et al. 2020; Roginski et al. 2024, Grandchamp et al. 2023, Grandchamps et al. 2024). Durand et al. (2019) addressed both problems simultaneously by characterizing the translation profile of closely related *S. paradoxus* populations, demonstrating a high turnover of translational activity in ORFs emerging *de novo* within species (Durand et al. 2019). This approach, combining genomic and translation data in a comparative framework, has shown great potential in deciphering the first steps of *de novo* gene birth at a short evolutionary timescale. Yet, it has so far been under-exploited, and has in particular never been implemented to assess the importance of short, translated intergenic ORFs in the *de novo* gene birth in plants.

Here, we aimed to characterize the early steps of *de novo* gene emergence in the *Arabidopsis* genus by specifically investigating the properties of intergenic ORFs (hereafter referred to as IGORFs) and annotated genes (referred to as canonical CDSs for coding DNA sequences) that have recently emerged *de novo*. We focused on *A. lyrata*, whose recent evolutionary divergence from its sister species *A. halleri*, and from the model *A. thaliana* provides high resolution to the synteny analysis (Tang et al. 2008). We produced ribosome profiling data from root and leaf tissue for *A. lyrata* and *A. halleri*, which we combined with publicly available data for *A. thaliana* to annotate signals of translation on both IGORFs and CDSs. We find that 142 IGORFs and 21 CDSs have emerged *de novo* along the *A. lyrata* lineage and show signs of ongoing translation. We observed that collectively, these translated *de novo* IGORFs and CDSs share many characteristics with untranslated intergenic ORFs: they are short, have a low GC content, prevalently use non-canonical START codons, and evolve in the absence of detectable selective pressure. Nonetheless, some of them possess characteristics closer to those of conserved genes, illustrating their potential functional relevance.

## Results

### Identifying intergenic ORFs and CDSs with translation signals

Our overall strategy was based on four closely related Brassicaceae species for which high-quality reference genome assemblies and protein-coding gene annotations were available: *A. lyrata* (Kolesnikova et al. 2023), *A. halleri* (Pavan et al. 2025), *A. thaliana* (Lamesch et al. 2012), and *Capsella rubella* (Slotte et al. 2013). Our main focal species, *A. lyrata,* diverged from *A. halleri* less than one million years ago, and from *A. thaliana* about five million years ago (Roux et al. 2011). We used Capsella as an outgroup genus, which has split from the other species around eight million years ago (Acarkan et al. 2000; Koch and Kiefer 2005). First, we annotated all possible ORFs in intergenic regions across these four genomes, using a relaxed sequence-based bioinformatic annotation of ORFs, allowing for alternative START codons (ATG, TTG, CTG, GTG, ACG), a minimal length of nine nucleotides, with no overlap with gene annotations (see Methods). This yielded a total of about 23, 19, 3 and 10 million IGORF annotations in *A. lyrata, A. halleri*, *A. thaliana* and *C. rubella,* respectively (**Table S1**). In comparison, 30,443, 34,805, 27,559 and 26,521 CDSs had been annotated as genes in these high-quality genome assemblies, respectively (**Table S1**). Second, we obtained experimental evidence for translation activity for the three Arabidopsis species. For *A. lyrata* and *A. halleri*, we grew plants under identical hydroponic conditions, extracted total RNAs from a controlled mixture of leaf and root tissue and produced new ribosome profiling data. After removing reads corresponding to residual rRNA contamination (up to 40%, see Methods and Table S2), we obtained 30 and 18 million mapped reads respectively for *A. lyrata* and *A. halleri*, which we used for the detection of translation signals. For *A. thaliana*, we took advantage of the high-quality ribosome profiling datasets from (Hsu et al. 2016; Wu and Hsu 2022), comprising a total of 40 to 110 million mapped reads from root, shoot and seedling tissues (**Table S2**).

We evaluated the quality of ribosome profiling libraries by examining metagene plots within the CDS of canonical protein-coding genes, and then annotated translation signals at individual IGORFs and CDSs. A high-quality ribosome profiling dataset would exhibit a noticeable nucleotide triplet periodicity, a stark increase in coverage at the start codon position, and a pronounced decrease at the stop codon position, reflecting the dynamics of ribosome occupancy along transcripts. The ribosome profiling dataset we produced for *A. lyrata* exhibited a strong periodicity signal, together with a sharp increase in coverage from the start position and a corresponding sharp decrease after the stop position (**Figure 1A**), indicating reliable evidence for translational activity. The signal of periodicity was especially apparent for 28-mers, which corresponds to the expected size of ribosomal footprints (**Figure 1A**, Ingolia et al. 2009).

**Figure 1:**
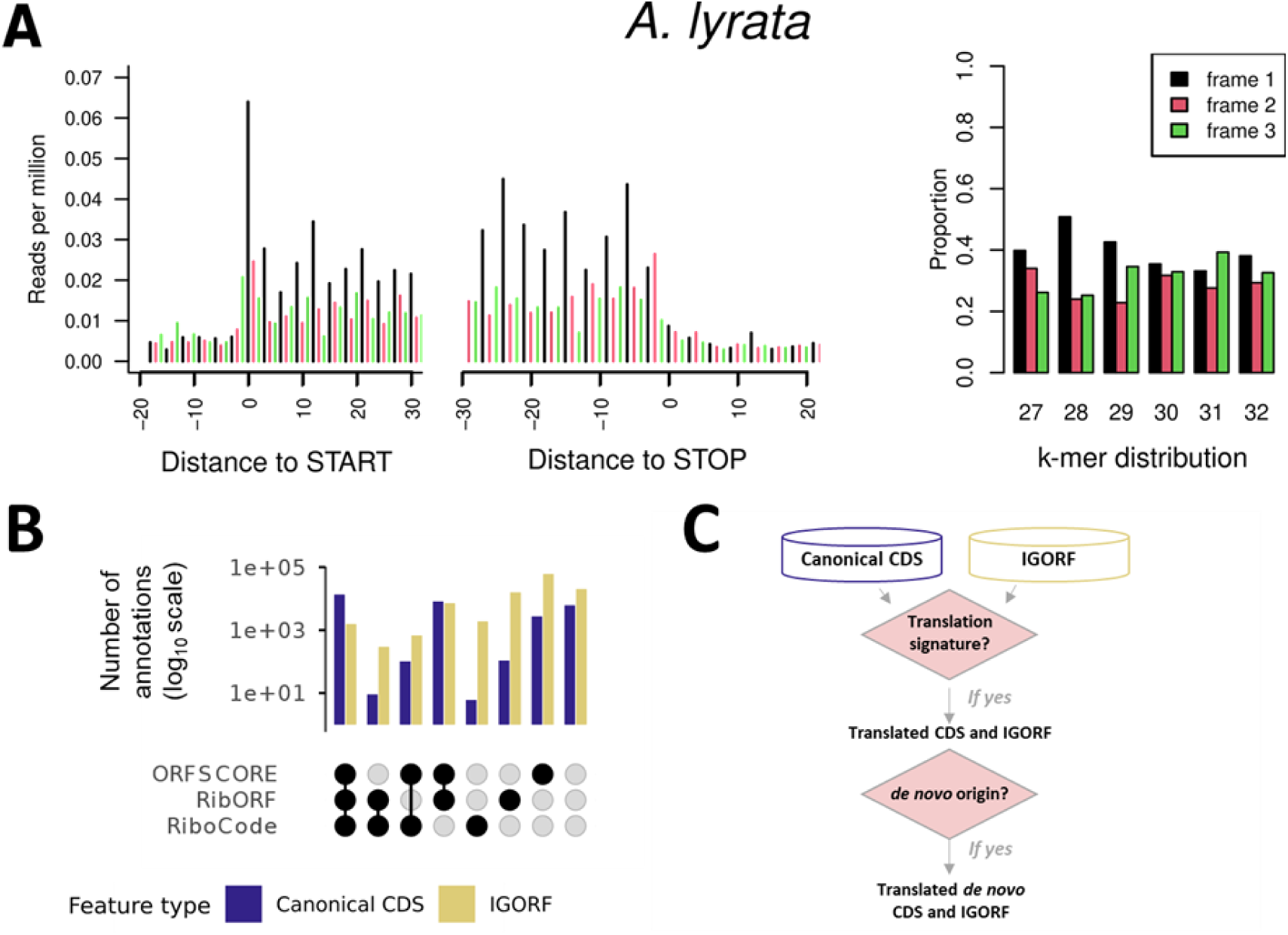
Detection of translated CDSs and intergenic ORFs (IGORFs) in *A. lyrata.* **(A)** Metagene plots of 28-kmers from ribosome profiling, after P-site shifting (reads were shifted according to their k-mer specific P-site offset, and trimmed to be 1 base-long), used for the annotation of translation events; and prevalence of k-mers depending on which frame they are found in, produced with modified RibORF scripts (ji 2018). Metagene plots are created by concatenating the per-nucleotide coverage at the P-site of ribosome profiling footprints within all known canonical protein-coding genes, measured in reads per million (RPM) relative to the start position of CDS. Coverage is colored based on its frame relative to the CDS: black for the first frame, red for the second frame and green for the third frame. The relative abundance of each k-mer, between 27 and 32 nucleotides in size, is also colored based on the frame in which the reads are located. Ribosomal footprints are expected to be around 28 nucleotides long (i.e. the length of the mRNA segment protected by the ribosome (Ingolia et al. 2009)). (B) Number of annotations detected as translated (log10) according to the methods that detected them as translated, and to the category to which they belong: canonical CDS or IGORFs. Black circles represent the detection of translation, grey circles represent its absence. (C) Graphic summary of the methodology to annotate translated de novo CDS and intergenic ORFs (see Figure S2 for the complete flowchart).

The identification of translation signals at individual IGORFs and CDSs remains a challenging task, especially in the context of the discovery of recently emerged *de novo* genes (Ruiz-Orera et al. 2014; Ruiz-Orera et al. 2018; Ruiz-Orera and Albà 2019; Papadopoulos et al. 2021). For this reason, we annotated translation signals using three independent methods. First, we used the least stringent ORFscore (Bazzini et al. 2014), which is based on the preferential accumulation of RPF reads in the first reading frame, taking the bias in canonical genes as a reference to distinguish true signal from noise, where reads are expected to accumulate across all reading frames uniformly. A second, more stringent method, RibORF (Ji 2018), uses Machine Learning trained on canonical genes to identify and recognize triplet periodicity. Finally, the third and most stringent method, RiboCode (Xiao et al. 2018), uses a modified Wilcoxon signed-rank test to annotate triplet periodicity *de novo.* To determine the accuracy of these annotations, we first examined CDS that were annotated as protein-coding genes. The majority of the CDSs annotated in *A. lyrata* (24,401 out of 30,443, or around 80%) were found to be translated by at least one of the three annotation methods (ORFscore, RibORF or RiboCode) (**Table S1**). Among them, 13,266 CDSs (44%) were found to be translated by all three methods (**Figure 1B, Table S1**). It should be noted that not all canonical CDSs are expected to be identified as being translated by this single experiment, due to inherent technical (*e.g.* assessing translation signals on lowly expressed regions) and biological limitations (*e.g.* limited number of sampled tissues, developmental stage-specific expression). Overall, we conclude that the signals we detect are powerful, but rather under-estimate the actual translation potential.

We then analysed the 23 million IGORFs annotated in *A. lyrata* (**Table S1**), and identified a small subset of 87,433 with detectable translation signatures by at least one method (**Figure 1B** and **Table S1**). Among those IGORFs with translation signatures, 1,548 displayed the most robust translation signals, *i.e.* they were identified by all three methods (**Figure 1B** and **Table S1**). Among the annotation methods we used, RiboCode stands out as the most stringent, with many IGORFs and CDSs identified as being translated by both ORFscore and RibORF. For downstream analyses, in an effort to be as exhaustive as possible, we focused on IGORFs and CDSs that were detected as translated by at least one method (**Figure 1C**).

For *A. halleri*, the ribosome profiling dataset displayed a less pronounced periodicity pattern compared to *A. lyrata*. Nevertheless, we were able to annotate 21,167 CDSs and 21,688 IGORFs with a translation signal detected with at least one method (**Figure S1**). For *A. thaliana*, the public dataset from Hsu et al. (2016) and Wu and Hsu (2022), optimized on the model species, had the best periodicity signal and allowed us to annotate 21,946 CDSs and 1396 IGORFs with translation signatures detected with at least one method (**Figure S1, Table S1**). Given the better quality of the ribosome profiling data from *A. lyrata* compared to *A. halleri*, for the following analyses we focused on the repertoire of IGORFs and CDSs with signals of translation detected in *A. lyrata*, and analysed their conservation and nucleotide sequence properties.

### Synteny analysis revealed translated IGORFs and CDSs that have emerged de novo in the Arabidopsis lineage

We then sought to identify which of the IGORFs and CDSs exhibiting translation activity in *A. lyrata* have appeared *de novo* in the Arabidopsis genus, *i.e.* after the divergence from *C. rubella,* which we used as an outgroup. To do so, we identified orthologous regions (genomic regions between two consecutive conserved protein-coding genes), which we aligned and filtered to obtain syntenic, orthologous regions of the translated IGORFs and CDSs in neighbouring species (see Methods, **Table S3, Figure S2**). Additional homology searches were performed to exclude sequences with homology to proteins outside of *Arabidopsis*, and/or outside of syntenic regions, thereby discarding orthology assignments lacking syntenic support **(Figure S2**). We annotated IGORFs and CDSs has having emerged *de novo* either because 1) they were fully absent (no coding sequence was present), 2) they contained a frameshift mutation or 3) they lacked a START and a STOP codon at the same positions, from the syntenic region in the outgroup (either *C. rubella,* or *A. thaliana* if the orthologous region was missing in *C. rubella* depending on the comparison).

Based on these stringent thresholds, we identified in *A. lyrata* a total of 142 translated IGORFs and 21 translated CDSs that have emerged *de novo* after the divergence from *C. rubella* (**Figure 2**; **Table 1**). Most of them had a clearly recognizable orthologous non-coding sequence in *C. rubella* and/or *A. thaliana* (**Figure 2**). 112 of the translated *de novo* IGORFs and six of the translated *de novo* CDSs were entirely specific to *A. lyrata* (*i.e.* they were not found either in the *A. halleri* nor in the *A. thaliana* genomes, Table 1). Similarly, 20 and 6 of *de novo* IGORFs and CDSs respectively were conserved in the *A. halleri* but not the *A. thaliana* genome, and may correspond either to gain in the ancestor of *A. halleri* and *A. lyrata* or to loss along the *A. thaliana* lineage. The IGORFs and CDSs identified as translated by two or three translation detection methods were either conserved or species-specific (**Figure 2**), which indicates that even recently emerged IGORFs and CDSs can already have acquired robust translation signals.

**Figure 2:**
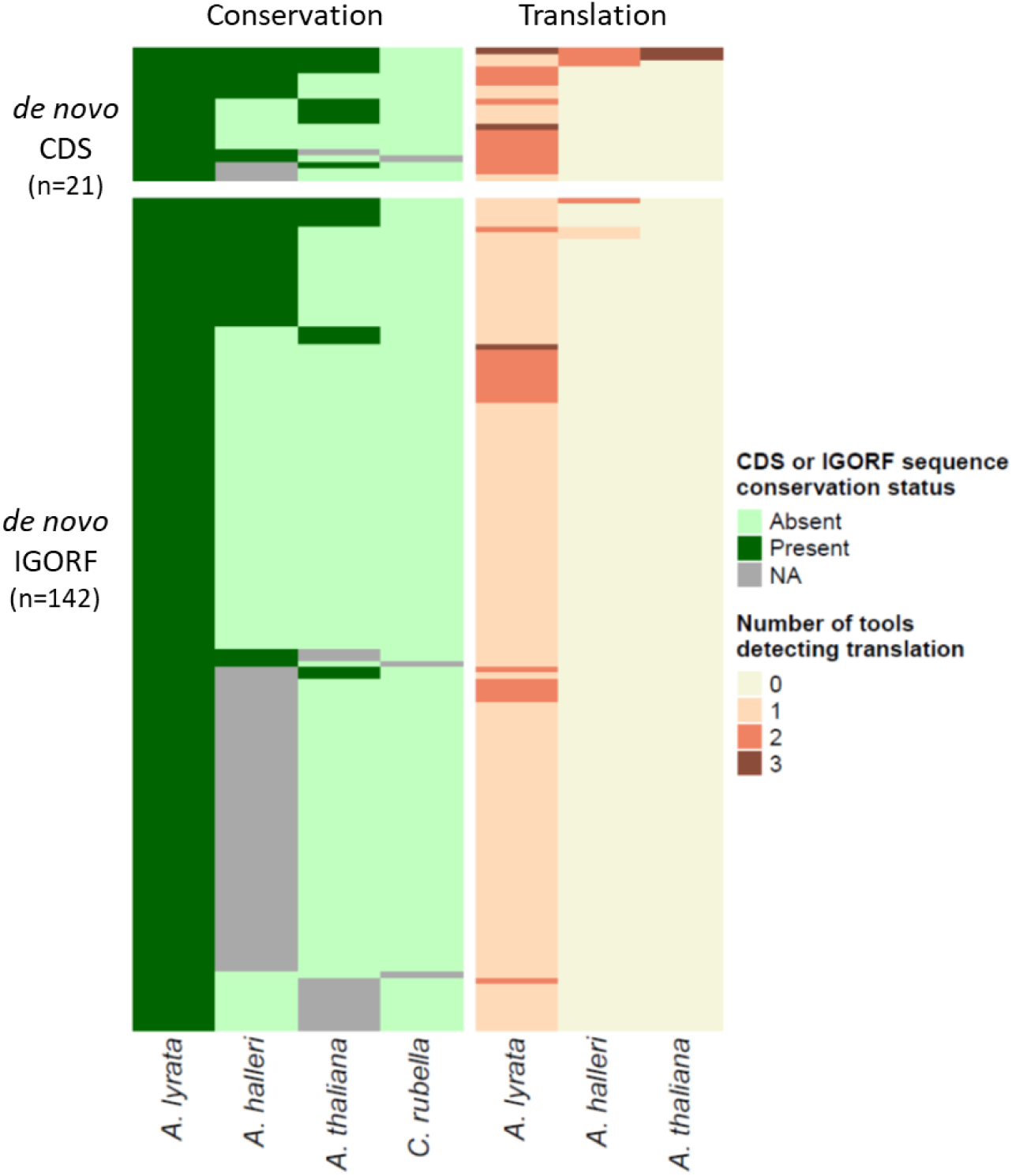
Heatmap of the conservation of the translated de novo IGORF and CDS sequences identified in A. lyrata (left), and translation detection (right). Each line represents an individual IGORF or CDS, and each column a species. Redundancy within each species was removed by keeping only the longest of overlapping de novo IGORF or CDS, and between species by removing duplicates of a same IGORF or CDS found to be translated in several species. A sequence was considered as conserved when its START and STOP positions, as well as its reading frame were conserved. If the IGORF or CDS is conserved in a species, it is annotated as “present”, if not it is annotated as “absent”. If no homologous syntenic region could be aligned, the IGORF or CDS was annotated as “NA”. As translation detection methods rarely annotate the exact same sequence positions as being translated, a given tool was considered as having detected translation if the exact IGORF or CDS position was detected as translated, or if an overlapping IGORF or CDS was detected as translated.

**Table 1:** Number of translated *de novo* CDS and IGORFs identified in *A. lyrata* by conservation category, and annotation type (*= already annotated as a CDS in at least one *Arabidopsis* species). Conservation categories are as follows: “Cons”, conserved in all *Arabidopsis* species; “Aly-Aha”, conserved in *A. halleri* and *A. lyrata* and “Aly”, *A. lyrata*-specific.

| Translated in: | Conservation category | *de novo* CDS* | *de novo* IGORF |
| --- | --- | --- | --- |
| *A. lyrata* | Cons | 9 | 10 |
|  | Aly-Aha | 6 | 20 |
|  | Aly | 6 | 112 |
|  | **TOTAL:** | **21** | **142** |

### Evolutionary trajectory of translated de novo IGORFs and CDSs

We then compared features of translated *de novo* IGORFs and CDSs to those of conserved orthologous CDSs on the one hand, and to those of untranslated IGORFs on the other hand. To do this, we defined orthologous CDSs as canonical CDSs with an identifiable ortholog in every species of interest and unt_IGORFs as IGORFs with no coverage by ribosome footprints. We classified *de novo* IGORFs and CDSs annotated in *A. lyrata* into successive conservation categories: conserved in all *Arabidopsis* species (hereafter “cons”), conserved between *A. halleri* and *A. lyrata* (“Aly-Aha”) and *A. lyrata-*specific (“Aly”) (**Table 1**, **Figure 3A**).

**Figure 3:**
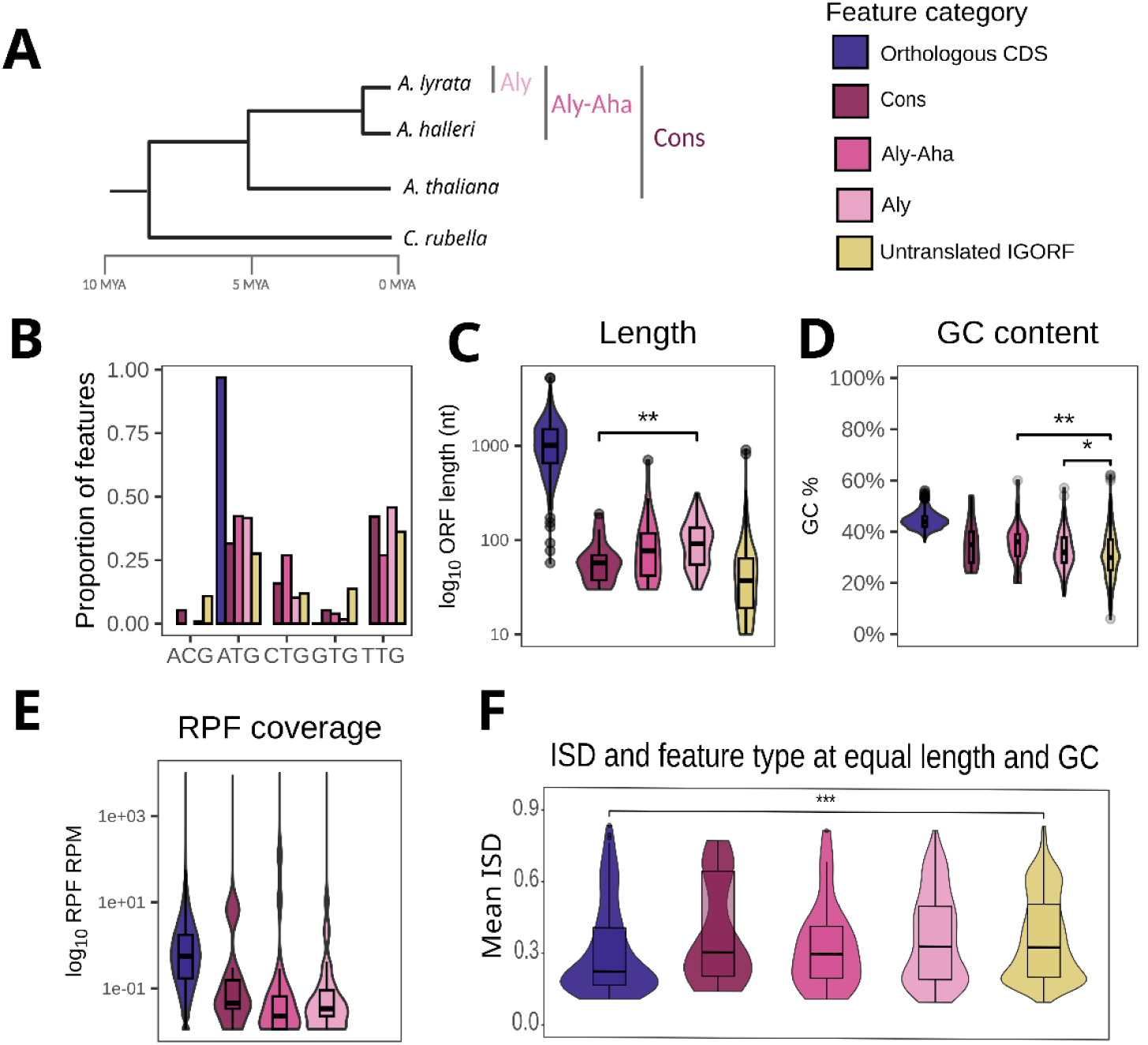
Characteristics of translated de novo CDSs and IGORFs compared to conserved canonical CDS and untranslated IGORFs in A. lyrata. De novo CDSs and IGORFs were classified according to their conservation category (A; see Table 1). For length and GC content (C,D), all statistically significant differences were represented, except for the ones between orthologous CDS and all other categories (Mann-Whitney U-test, * = p < 0.05, ** = p < 0.01, *** = p< 0.001).

First, we examined usage of the canonical (ATG) and alternative (ACG, CTG, GTG, TTG) start codons of the genetic code. As gene prediction methods generally consider canonical start codons only, conserved orthologous CDSs almost exclusively start with ATGs (**Figure 3B**). In *de novo* IGORFs/CDSs, the TTG start codon was equally prevalent as the canonical ATG, and other alternative start codons were also common, representing up to 25% of all *de novo* IGORFs/CDSs. These patterns were also observed across conservation classes of unt_IGORFs (**Figure S3**), and hence reflect the nucleotide composition of these sequences. Second, we observed that, as expected, conserved orthologous CDSs were on average longer (mean length = 1,111 nt) than unt_IGORFs (mean length= 52 nt), while *de novo* IGORFs/CDSs were intermediate (mean length = 100 nt) (**Figure 3C**). *De novo* IGORFs/CDSs that are conserved in the three Arabidopsis species (cons) were significantly shorter (mean length = 66 nt) than those that are specific to *A. lyrata* (Aly; mean length = 104 nt; p-value = 2.6 × 10^-3^). A likely reason for the length difference is the stringent way we defined conservation, as longer IGORFs and CDSs are more likely to be interrupted by mutations compared to shorter ones, and hence constitute larger mutational targets (Iyengar and Bornberg-Bauer 2023). Unt_IGORFs followed the same trend across conservation classes (**Figure S3**), lending support to this interpretation . Third, we observed a continuous decrease of GC content, from 44% in conserved orthologous CDSs, through a significantly lower 35% in the *de novo* IGORFs/CDSs that are *A. lyrata*-specific (Aly conservation class), down to a median of 30% for unt_IGORFs. The Aly-Aha and Aly *de novo* IGORFs/CDSs conservation classes had a lower GC content than conserved orthologous CDSs, but a significantly higher GC content than unt_IGORFs (**Figure 3D**). In contrast, no variation of GC content was observed when comparing unt_IGORFs of different conservation classes (**Figure S3**). When comparing *de novo* IGORFs/CDSs and unt_IGORFs belonging to the same conservation classes, we observe that the *A. lyrata*-specific *de novo* IGORFs/CDSs have a statistically significantly higher GC content than their unt_IGORFs counterparts (**Figure S4**). Fourth, in line with previous observations (Carvunis et al. 2012), translation activity (quantified by ribosome profiling reads per million, RPM) of the *de novo* IGORFs/CDSs was lower than that of conserved orthologous CDS (on average between 1.29 RPM versus 2.43 RPM, **Figure 3E**). Still, the two distributions overlap largely, and a substantial fraction of the *de novo* IGORFs/CDSs exhibit a level of translation activity that is comparable to that of many conserved orthologous CDSs (**Figure 3E**). To verify that the overall difference in translation activity was not simply caused by the difference in length of the annotations, we used a linear least squares regression, which showed that length alone could explain only about 2.6% of the observed variance in read depth (adjusted R-squared=0.02619, *p-value*<2.2 × 10^-16^). Furthermore, the differences in translation activity were also apparent when length differences were controlled for by calculating read depth only within the first 60 nucleotides of each feature, where the translation signal is most pronounced. Using a linear least squares regression, there was no evidence that length could explain the observed variance in read depth within the first 60 nucleotides (adjusted R-squared=-0.0001479, *p-value*=0.3675).

Next, we examined the levels of intrinsic structural disorder (ISD) of the predicted polypeptides. We predicted ISD using AIUPred (Erdős et al. 2021) on the *de novo* IGORFs/CDSs, as well on a random sub-sample of 1,000 orthologous CDSs and 1,000 unt_IGORFs for comparison (**Figure S5**). Taken at face value, our results indicate that unt_IGORFs and *de novo* IGORFs/CDSs exhibit significantly higher levels of ISD than conserved orthologous CDS (Mann-Whitney U test, *p-value*<2.2 × 10^-16^, **Figure S5**). However, previous studies have highlighted that polypeptides that are short have a lower propensity to form structural elements less readily, and thus tend to exhibit higher ISD than longer ones (Dill 1990; Peng et al. 2006). A similar pattern has been reported for polypeptides encoded by GC-rich nucleotide sequences and their higher chance of triplets encoding disorder-promoting amino acids (Basile et al. 2017). These two technical factors are known to affect the accuracy of disorder prediction algorithms (Peng et al. 2006). Indeed, across unt_IGORFs and conserved orthologous CDSs, disorder decreases with length (unt_IGORFs: Spearman *⍴(disorder,length*)= -0.24, *p-value*= 1.22 × 10^-14^; orthologous CDSs: *⍴(disorder,length*)= -0.14, *p-value*=1.29 × 10^-5^), and increases with GC content (unt_IGORFs: Spearman *⍴(disorder,GC*)= 0.13, *p-value*= 2.53 × 10^-5^; orthologous CDSs: *⍴(disorder,GC*)= 0.15, *p-value*=1.67 × 10^-6^, **Figure S5**). Yet, both within and across translated *de novo* IGORFs/CDSs conservation categories, no such significant relationship could be established between disorder and length or GC, with the exception of the *A. lyrata-*specific conservation class (Aly), where GC content was unexpectedly negatively correlated to ISD (*⍴(disorder,GC*)= -0.20, *p-value*=0.01474). A modest but significant proportion of the variance in ISD is jointly explained by sequence length and GC content (adjusted R² = 0.0353, *p* < 2.2 × 10^-16^ for length; adjusted R² = 0.0844, *p* < 2.2 × 10^-16^ for GC content, **Figure S5**). Together, these results confirm a confounding effect of GC content and protein length on ISD. To control for these two factors jointly, we predicted ISD for all conserved orthologous CDS and kept only those matching the length and GC content of unt_IGORFs and *de novo* IGORFs/CDSs (<139 amino acid residues long, 21-39% GC content). Among the 160 conserved orthologous CDSs that matched these stringent criteria, we still observed a statistically significant difference in ISD between categories (**Figure 3F**). Hence, we conclude that *de novo* IGORFs/CDSs and unt_IGORFs appear to be more disordered than conserved orthologous CDS even after controlling for differences in length and GC contents.

### A select few de novo IGORFs and CDSs have the potential to be proto-genes

Out of the 163 translated *de novo* IGORFs/CDSs which we identified, a few stand out as already having notable gene-like characteristics. Indeed, 21 had already been annotated as CDSs (**Table 1**). Among them, the six CDSs that are specific to *A. lyrata* contain one or no intron, which is consistent with the low number of introns generally observed in young *de novo* genes (Zhang et al. 2019). For example, the *A lyrata*-specific d*e novo* CDS annotated as AL6G18580 (**Figure 4A**) has no sequence homology to any previously described protein in any species outside *A. lyrata*, displays translation signals by two methods (ORFSCORE and RibORF), starts with the canonical start codon ATG, and is 222 nucleotides long (73 amino acids). An orthologous sequence was present in *C. rubella,* where it was clearly non-coding, with no conservation of the START and STOP codon and of the frame. While the START and STOP codon sequences are conserved in *A. halleri* and *A. thaliana*, a few indels interrupt the reading frame in these species. Hence, this particular *de novo* CDS could have emerged either specifically in *A. lyrata* or in a more ancestral *Arabidopsis* species, and have been lost secondarily in *A. thaliana* and *A. halleri* by indels. Supplementary BLAST alignments (**Table S4**) revealed that this locus had previously been predicted as a non-coding RNA in *A. lyrata* (Hupalo and Kern 2013). Further alignments (**Figure S6**) revealed that AL6G18580 is homologous to the upstream region of the *A. thaliana* miR162a miRNA, encompassing the start of the precursor and the future mature miRNA. Given the high conservation degree of miR162a (conserved beyond *Brassicaceae*), the upstream region of the miR162a locus may have recently acquired a translated *de novo* CDS in *A. lyrata*. This is reminiscent of the discovery of polypeptide-encoding ORFs within the 5’ region of miRNA primary transcripts in several plant species, including *A. thaliana* (Lauressergues et al. 2015; Lauressergues et al. 2022). AlphaFold2 confidently predicts that the potential encoded polypeptide forms an alpha-helix (predicted local distance difference test (pLDDT) between 70 and 90), facing five beta-sheets with varying prediction confidence (from pLDDT < 50 to >90, **Figure 4A**).

**Figure 4:**
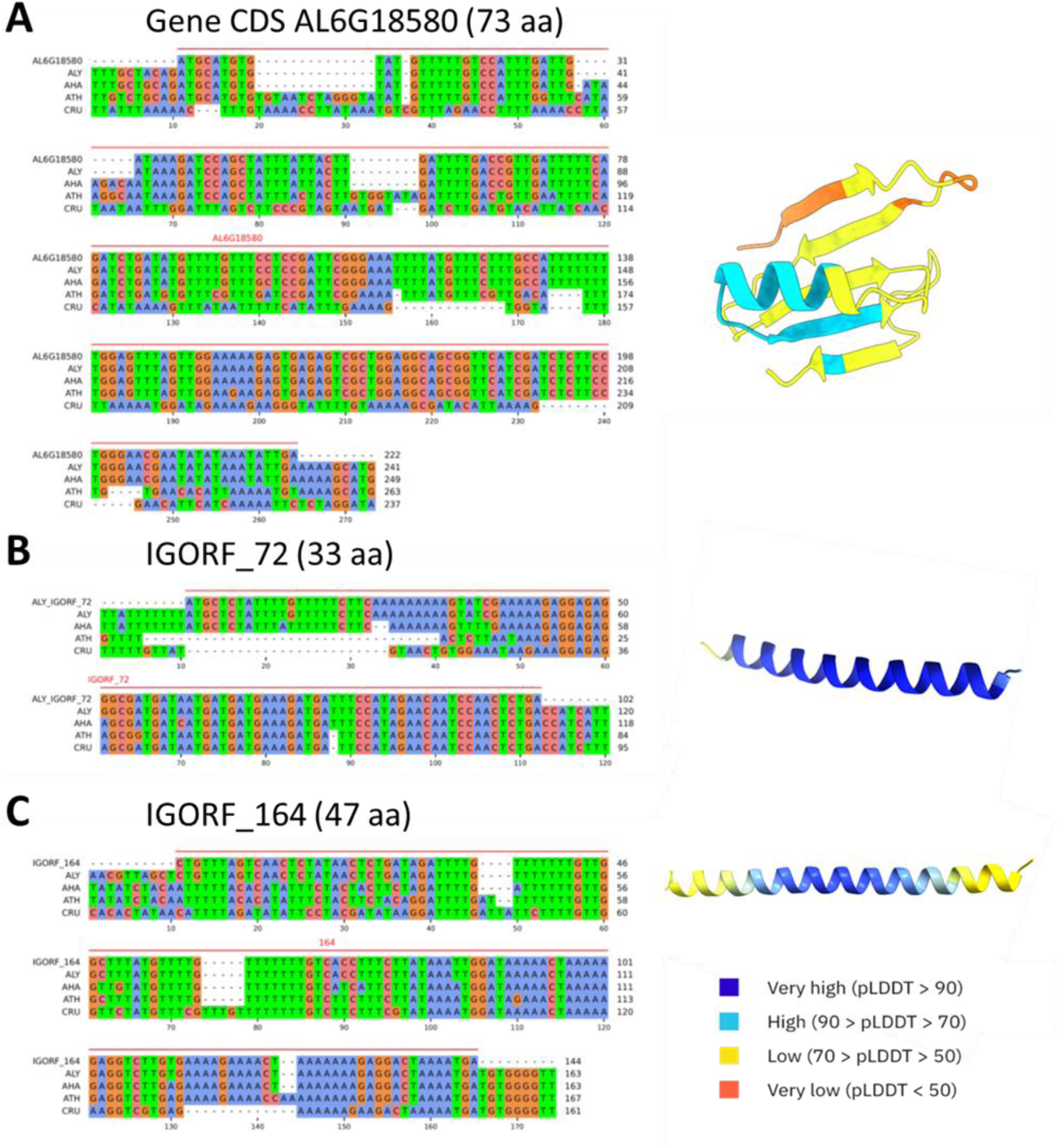
Alignments of a selection of de novo gene candidates specific to A. lyrata (CDS AL6G18850, A; IGORF_72, B and IGORF_164, C) with their orthologous genomic intergenic regions in the other species. For each alignment, the first line is the sequence of the CDS or the IGORF, the next four lines represent the orthologous genomic regions for each species: ALY for A. lyrata, AHA for A. halleri, ATH for A. thaliana and CRU for C. rubella, extended by 10 nucleotides on both sides. Alignments are colored by nucleotides and visualized using pyMSAviz (Shimoyama 2022). The CDS or IGORF itself is represented by a red line above the alignments. The corresponding AlphaFold2 structure prediction of the encoded polypeptides is displayed next to each corresponding alignment. Residues are colored by pLDDT score (see legend).

Beside such *de novo* CDSs, most of the translated *de novo* sequences we identified were actually intergenic (142 IGORFs, **Table 1**), and some of them have features similar to the *de novo* CDSs. Here, we highlight two examples of *A. lyrata*-specific IGORFs: IGORF_72 and IGORF_164. The IGORF_72 **(Figure 4B**) was detected as translated by two methods (ORFSCORE and RibORF) in *A. lyrata*, is 102 nucleotides long (33 amino acids) and starts with a canonical start codon (ATG). The structure of the corresponding potential encoded polypeptides is predicted to be a single alpha-helix with very high confidence (pLDDT > 90, **Figure 4B).** The IGORF_164 (**Figure 4C**) was detected as translated by one method (RibORF) in *A. lyrata*, is 144 nucleotides long (58 amino acids) and starts with an alternative start codon (CTG). The predicted structure of the polypeptide consists of a single alpha-helix, with very high confidence in the central region (pLDDT > 90) that gradually decreases toward both extremities (pLDDT 50–70; **Figure 4C**).

For both IGORFs examples, the orthologous sequences exhibit indels or single nucleotide polymorphism (SNP) in every other species we studied, leading to the loss of a STOP codon in *A. lyrata*, resulting in an expansion of the IGORF size. The IGORF sequences exhibit no detectable homology to any known protein-coding gene or non-coding locus (see Methods, Figure S2 and Table S5).

### The repertoires of de novo IGORFs and CDSs are highly labile within species

To gain insight into the conservation of the translated *de novo* IGORFs/CDSs at a finer phylogenetic scale, we then analyzed five high-quality genome assemblies from natural accessions of *A. lyrata* that we either collected from the literature (Padilla-García et al. 2025; Scott et al. 2025) or assembled *de novo*. We identified syntenic regions based on the transfer of gene annotations from the *A. lyrata* reference accession (MN47, Kolesnikova et al. 2023). The conservation of each *de novo* IGORF or CDS within syntenic regions that were conserved between accessions was determined using the previous filters and alignments (**Figure S2**). Even within *A. lyrata* species, a high degree of lability was apparent (**Figure 5A**). Specifically, very few of the *de novo* IGORFs/CDSs were conserved across all five accessions, with 21% being accession-specific, and only 29% found in more than two accessions. We then compared these patterns of conservation to those of unt_IGORF, which we used as a neutral control. The *de novo* IGORFs/CDSs were not more conserved across accessions than unt_IGORFs in *A. lyrata* (permutation test, *p*-value=0.442), and we conclude that the repertoires of *de novo* IGORFs/CDSs are highly labile within species.

**Figure 5:**
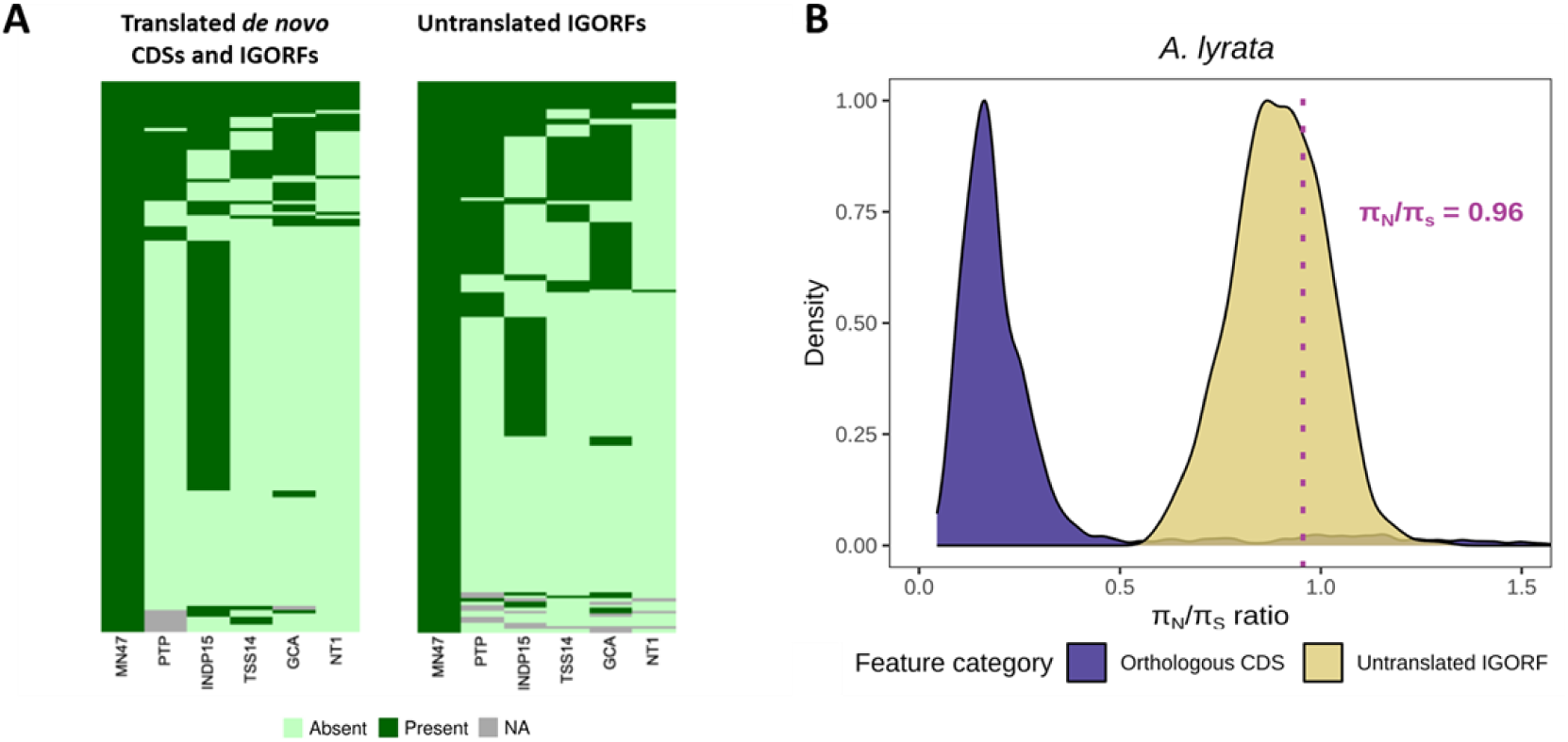
A) Heatmap of the conservation of translated de novo IGORFs/CDSs and untranslated IGORF (unt_IGORF) sequences across accessions from A. lyrata . Each line represents an individual IGORF or CDS, and each column an accession. NA indicates that no homologous syntenic region could be aligned. Only IGORFs/CDSs present in the reference accession MN47 and with an orthologous genomic region detected in at least one additional accession were included (n = 152). B) The πN/πS ratio of de novo IGORFs/CDSs (dotted vertical line; a single value computed after concatenation) matches that of untranslated intergenic ORFs (random subset of 1,000 ORFs) and departs from that of conserved canonical CDSs (random subset of 1,000 CDSs), indicating that the strength of purifying selection on them is either null or at most very weak .

### No evidence for purifying selection on de novo IGORFs and CDSs

Finally, we sought to evaluate the strength of purifying selection acting on translated *de novo* IGORFs/CDSs. To do this, we collected genomic resequencing data from a range of natural populations of *A. lyrata* (a total of 39 individuals) (**Table S5**). We first evaluated total nucleotide diversity per site, and found that conserved orthologous CDSs display the lowest levels of diversity (on average 5.60 × 10^-5^), and unt_IGORFs the highest (on average 1.67 × 10^-4^) (**Figure S7**). *De novo* IGORFs/CDSs displayed levels of nucleotide diversity similar to those of unt_IGORFs, with an average of 9.12 × 10^-5^ (*p*=0.39), and much higher than those at conserved orthologous CDSs (*p*=3.6 × 10^-6^; **Figure S7**). The ratio of non-synonymous (*π*_N_) to synonymous nucleotide diversity (*π*_S_) revealed strong purifying selection on conserved orthologous CDSs (*π*_N_/*π*_S_=0.32). The average ratio for untranslated IGORFs was close to one (0.90), corresponding to the expectation under neutral evolution (**Figure 5B**). Because d*e novo* IGORFs/CDSs tend to be short, we concatenated their sequences to get more polymorphism in order to be able to perform the *π*_N_/*π*_S_ ratio analysis. Translated *de novo* IGORFs/CDSs exhibited a *π*_N_/*π*_S_ ratio of 0.95 in *A. lyrata*, landing them right in the distribution of unt_IGORFs *(p*=0.57). Hence, the strength of purifying selection on *de novo* IGORFs/CDSs taken as a group does not detectably differ from that on our neutral control (unt_IGORFs), and is thus either null or at most very weak.

## Discussion

### Pervasive translation of intergenic ORFs in Arabidopsis

Our study revealed that translation of IGORFs is common in *Arabidopsis* species, even when stringently restricting the analysis to those with the highest evidence of translation. The numbers of IGORFs with translation signals we observed in *A. lyrata* (1,548 with the most robust translation signals) are consistent with previous reports in *A. thaliana*, where 2,996 ORFs were found to be expressed in intergenic regions (Hanada et al. 2007) and 2,099 highly expressed under certain experimental conditions (Hanada et al. 2013). They are also consistent with reports across distant clades such as the yeast *S. cerevisiae*, where 1,266 IGORFs were either occasionally or highly translated (Papadopoulos et al. 2021), the yeast *S. paradoxus*, where 447 IGORFs were translated (Durand et al. 2019), the mouse *M. musculus*, where 1,980 non-conserved ORFs were translated (Ruiz-Orera et al. 2018), or the fly *D. melanogaster*, where 343 small ORFs were found to be translated (Patraquim et al. 2020).

While it is clear that translation is common in intergenic regions, the exhaustive characterization of such non-canonical translation events remains a major challenge. Indeed, ribosome profiling may introduce biases, both during laboratory experiments and computational analyses (Bartholomäus et al. 2016). For example, translation elongation inhibitors do not homogeneously stop ribosomes along transcripts and may be affected by codon composition (Orelle et al. 2013), making it more difficult to pin-point the exact positions at which translation is happening (Bartholomäus et al. 2016). This makes the precise identification of which ORFs are actively being translated difficult, especially in non-coding regions, and as we observed, different analysis methods can differ sensibly in their predictions. Dosage of RNAse I is delicate and of crucial importance, and we therefore relied on the recommended dosage for *A. thaliana* (Wu and Hsu 2022). Still, we observed high levels of rRNA contamination (up to 40%), which might have partly been the product of too high of a RNAse I dosage leading to ribosome degradation (Becker et al. 2013; Bartholomäus et al. 2016), or to inefficiency of the experimental ribosomal depletion, with varying results across samples. These biases affect all coding sequences, but render non-canonical translation events particularly difficult to detect, where different prediction methods differ the most, and ORFs are not saturated in coverage at usual sequencing depths (Lei et al. 2024). All of these factors contribute to diminishing the statistical power to uncover translation events outside of genic regions. To minimise these biases and maximize the chances of detecting translation events, a recommended strategy is to combine datasets (Wacholder et al. 2023). Since no other public datasets were available for *A. lyrata* and *A. halleri,* we instead maximized sequencing depths, as recommended by Lei et al. (2024), reaching up to 639 million reads (as opposed to the typical range of 150 to 200 million reads in available *A. thaliana* ribosome profiling datasets (Hsu et al. 2016; Wu and Hsu 2022). Finally, it has previously been reported that non-canonical translation, including that of *de novo* genes, tends to happen in a tissue-specific manner (Wu et al. 2014; McLysaght and Hurst 2016). For instance, in *A. thaliana*, the *de novo* gene *SWK* was found to be highly expressed specifically in seeds and siliques (Jin et al. 2025). Therefore, an exciting next step will be to determine the extent to which the non-canonical translation signals we detected vary across developmental stages, across plants collected from diverse environments and across a broad panel of genetic origins. Overall, beyond these variations and inherent technical limitations, it is clear from our results that translation activity is not restricted to canonical genic sequences in Arabidopsis species, as is now recognized in a range of model and non-model organisms (Ingolia et al. 2014; Ruiz-Orera et al. 2018; Durand et al. 2019; Chen et al. 2020; Sruthi et al. 2022; Papadopoulos et al. 2024).

### Frequent emergence of de novo IGORFs and CDSs

Such pervasive transcription (Neme and Tautz 2016; Schmitz et al. 2018) and translation (Ingolia et al. 2014; Ruiz-Orera et al. 2018; Blevins et al. 2021; Wacholder et al. 2023; Papadopoulos et al. 2024) has previously been described as the “transient translatome” (Wacholder et al. 2023). The presence of numerous species-specific translated IGORFs (112 in *A. lyrata*) supports the idea of frequent gains of transcription and translation in regions considered to be “non-coding”, which would become temporary, initial substrates for the *de novo* emergence of genes (Schmitz et al. 2018; Durand et al. 2019; Grandchamp et al. 2024). Phylogenetic conservation of these translated *de novo* IGORFs is rather low, both across species, or across accessions within species, and their levels of nucleotide diversity and *π*_N_/*π*_S_ ratio indicate that they globally evolve neutrally, in line with *e.g*. Ruiz-Orera et al. (2018).

### The first steps of de novo gene emergence

The processes by which some *de novo* IGORFs and CDSs deviate from noise and eventually acquire gene-like characteristics remain somewhat unclear. Here we found that many of them show similarity with untranslated intergenic ORFs: they have a shorter length than conserved orthologous CDSs, a high prevalence of non-canonical START codons, are overall translated at lower levels and collectively exhibit no sign of purifying selection, which corresponds to what has previously been observed in yeast, mice and fruit flies (Carvunis et al. 2012; Ruiz-Orera et al. 2018; Vakirlis et al. 2018; Papadopoulos et al. 2021; Grandchamp et al. 2023; Wacholder et al. 2023). Strikingly, the GC content of *de novo* IGORFs/CDSs was intermediate between that of conserved orthologous CDS and that of untranslated IGORFs, and increased continuously with phylogenetic conservation. A similar continuum of GC content was observed in *A. thaliana* and *Drosophila* (Li et al. 2016; Peng and Zhao 2024), but a different trend was observed in yeast, where *de novo* genes tend to emerge in regions that are richer in GC content than in conserved CDS (Vakirlis et al. 2018) and in humans, where *de novo* genes can arise from long non-coding RNAs (lncRNAs) exhibiting significantly higher GC content than protein-coding genes globally (An et al. 2023). A recent study has highlighted that GC-rich IGORFs are more likely to give rise to novel genes *de novo*, thanks to the greater stability and expression level of GC-rich RNAs (Roginski et al. 2026). GC-content has also been linked to the foldability and solubility of the encoded polypeptides, and intermediate levels of GC-content are associated with greater foldability, with older *de novo* genes showcasing greater foldability than younger ones across a wide range of taxa (Bornberg-Bauer and Eicholt 2026; Roginski et al. 2026). As the number of species where the emergence of *de novo* genes is studied increases in the future, it will be exciting to finally have the possibility to test whether the trends observed across species can be generalized as global mechanisms for *de novo* gene emergence.

GC content is also strongly correlated to other molecular features, including ISD (Basile et al. 2017; Nielly-Thibault and Landry 2019). Similarly to what we observed in conserved orthologous CDSs and unt_IGORFs, a positive relationship between GC content and disorder, and a negative one between length and disorder have been documented in the literature (Peng et al. 2006; Basile et al. 2017). Short length in general, regardless of conservation or expression, has been reported to be associated with higher disorder (Middendorf and Eicholt 2024). In non-conserved *de novo* genes in *Drosophila*, higher GC content and shorter length are both contributing factors to the observed increase in disorder compared to other gene categories (Zheng and Zhao 2022; Middendorf et al. 2024). In translated non-canonical ORFs from humans and mice, a similar pattern of higher disorder linked to shorter length has been documented (Schmitz et al. 2018). While the shorter length of *de novo* IGORFs/CDSs in our data may contribute to their higher disorder, their lower GC content should act in the opposite direction, contributing to lower disorder. Furthermore, in species-specific *de novo* IGORFs/CDSs, we observe a significant negative relationship between GC and disorder. Keeping in mind that analysing short sequences is challenging, our analyses suggest that translated *de novo* IGORFs and CDSs are more disordered than conserved orthologous CDSs, even when correcting for their different length and GC content. This finding is in line with previous observations in *Drosophila* (Middendorf et al. 2024).

### A few translated de novo IGORFs and CDSs are promising gene candidates

While most *de novo* IGORFs/CDSs possess features similar to those of unt_IGORFs, a few of them are closer to the genic end of the spectrum and are promising *de novo* gene candidates. In fact, a small subset of those we identified had already been annotated as proper CDSs (21 in *A. lyrata*). Several of them display unambiguous evidence of active translation, and are predicted to have the potential to adopt simple 3D structures such as alpha helices or beta sheets, in line with structure predictions of *de novo* genes in other species (Chen et al. 2024; Middendorf et al. 2024; Peng and Zhao 2024). Having established that these CDSs actually correspond to *de novo* genes opens the exciting possibility to explore their functional properties in more detail, especially given the wealth of molecular resources available in the model *A. thaliana*. In addition to those that had already been spotted as CDSs, our analyses revealed a substantial number of previously unannotated intergenic ORFs displaying similarly strong evidence of translation, with similar length and structure predictions as the *de novo* CDSs. While previous plant studies have mostly focused on the functional potential of previously annotated *de novo* emerged genes with already well-established gene-like features (Zhang et al. 2019; Jin et al. 2025), our results highlight the value of investigating the properties of intergenic translated ORFs as raw material for the emergence of new genes *de novo*.

Overall, our findings demonstrate that pervasive translation takes place even in intergenic regions in *Arabidopsis* species, providing a vast reservoir of transiently, lowly expressed, mostly neutrally evolving ORFs. Among those, a select few may ultimately give rise to functional genes, by progressively becoming longer, more highly expressed, and acquiring a higher GC content. Future work combining ribosome profiling, mass spectrometry, and functional assays across a variety of tissues and developmental stages will be essential to validate this repertoire of *de novo* peptides and determine their physiological relevance, particularly how they interact with other proteins and potentially integrate into molecular interaction networks.

## Material and methods

### Plant Material

Three seeds of either *A.halleri, A. lyrata* or *C. rubella* were placed on each of ten 0.7% agar-filled micro-centrifuge tub lid and imbibed in germination medium as described in (Conn et al. 2013), then left in the dark at 4°C for two days. They were then transferred to hydroponic growth tanks under standard greenhouse conditions, where the seeds germinated and seedlings were grown hydroponically in a standard growth medium composed of 1 mM Ca(NO3)2, 0.5 mM MgSO4, 3 mM KNO3, 0.5 mM NH4H2PO4, 0.1 μM CuSO4, 0.1 μM NaCl, 1 μM KCl, 2μM MnSO4, 25 μM H3BO3, 0.1 μM (NH4)6Mo7O24, 20 μM FeEDDHA, and 1 μM (for *A. lyrata*) or 10 μM (for *A. halleri*) ZnSO4. The pH of the solution was maintained at 5.0 using MES acid buffer (2 mM), and was replaced every week. Leaves and roots were collected and flash frozen in liquid nitrogen progressively along the growth of each plant over the course of three months (July to September) for one individual per species, and stored at -80°C.

### Preparation of RNA lysates

One ribosome profiling library was prepared for *A. lyrata* and one for *A. halleri*. We pooled each tissue sample collected on different dates to reduce the effect of the day of sampling and of the environment, as well as to ensure that we have sufficient material for the experiment. Tissue types were separately ground into a powder using liquid nitrogen and a mortar and pestle. We then suspended 0.25 g of each tissue powder in 4 mL of lysis buffer, as described in (Wu and Hsu 2022). The lysates were vortexed twice, transferred into 2 mL tubes and then agitated at 4°C for 10 min. All lysates were then spun at 5 000 × g at 4°C for 3 min; and the supernatant was aliquoted into refrigerated 1.5 mL tubes. The aliquots were spun at 20 000 × g at 4°C for 10 min and the supernatants were transferred to refrigerated 1.5 mL tubes. Aliquots of 200 μL were stored at -80°C and a single 10 μL aliquot was kept for the assessment of RNA concentrations by a Qubit RNA HS assay (Thermo Fisher Scientific). As it was apparent that root aliquots contained on average lower RNA concentrations than those from leaves, root and leaf aliquots were pooled per individual so as to control for 75% of RNA from roots and 25% from leaves. In the end, we have produced three lysate replicates per species.

### Ribosome footprints

The lysates were treated with 50 units RNase I Ambion (Invitrogen, AM2295) per 40 μg of RNA and digested for one hour on a nutator at 25°C. The samples were then moved to ice and 30 μL of SUPERase-IN (Fisher scientific, 10773267) were added to interrupt digestion. A sucrose cushion solution was prepared as described in (Baudin-Baillieu et al. 2016) and digested lysates were deposited on the cushions and centrifuged at 4°C and 657,000 g in a TLA110 rotor for 90 min. Pellets were then washed twice with 500 μL lysis buffer and resuspended in 750 μL lysis buffer; after which they were solubilized by pipetting up and down and transferred in a 2 mL microtube. Total RNA was extracted using a standard hot phenol procedure as described in (Baudin-Baillieu et al. 2016). A 17% polyacrylamide, 7M TAE-urea gel was pre-migrated at 100 V for 30 mins and the totality of extracted RNAs from each aliquot was charged and migrated for 5 h with size markers at 28 and 30 nucleotides. After migration, gels were stained with SYBR-GOLD and gel slices between the 28 and 30 nt markers were excised. The excised gel band was pulverized and eluted in 500 uL of extraction buffer (0,3 M CH_3_COONa pH 5,5, 1 mM EDTA) and then spun at 10 rotations/minute at 4°C overnight. The gel residues were retained on a Spin-X cellulose acetate filter 0,45 *μ*m Corning column (Dutscher; 2515806); and centrifuged at maximal speed for 5 mn. Per tube, 2*μ*L of glycogen (ThermoFisher; R0551) were added with 2 to 2,5 volumes of 100% EtOH and were then precipitated overnight at -20°C. Tubes were then centrifuged at +4°C at 21,000 g, and the pellets were dissolved in 15 μL of distilled water. The final RNA concentration was measured by fluorescence with the micro RNA Quant-iT kit (Thermofisher Scientific).

### Library preparation and sequencing

Samples were depleted in ribosomal RNAs according to the depletion protocol of the TruSeq® Stranded Total RNA Library Prep Plant kit (Illumina, 20020610). The obtained footprints were subsequently purified according to the following protocol modified from the Zymo RNA clean and concentrator kit (Zymo Research; R1017). An RNA binding buffer was adjusted with an equal, 100% ethanol volume, and two volumes (180 µL) of this adjusted buffer were added to the rRNA-depleted ribosome profiling samples (90 µL). Then, 1 volume of 100% ethanol (270 µL) was added to the mix to favor the retrieval of small RNAs between 17 and 200 nucleotides (ribosome footprints are typically approximately 30 nucleotides long). The total volume of 540 µL was deposited on a column, and centrifuged for 30s at 16,000 x g at room temperature. Then, 400 µL of RNA Prep Buffer were added to the column, which was again centrifuged under identical conditions, followed by the addition of 700 µL of RNA Wash Buffer and a third identical centrifugation. Finally, 400 µL of RNA Wash Buffer were added and the column was centrifuged for 1 min at 16,000g at room temperature. The column was put on a 2mL tube and the ribosome footprints were eluded on the column with 15 µL of distilled, nuclease-free water and centrifuged for 1 min at 16,000g at room temperature again. The final RNA concentration was measured by fluorescence with the micro-RNA Quant-iT kit (Thermofisher Scientific; Q32882).

The sequencing libraries were prepared according to the Biolabs NEBNext Multiplex Small RNA library Prep kit’s standard protocol with 12 PCR cycles. The resulting libraries were purified with the Monarch PCR & cleanup kit (Biolabs) and concentrations were assessed with the Quant-iT DNA HS kit. Sequencing was performed with a NextSeq system by the I2BC High-throughput sequencing facility, Gif-sur-Yvette.

### ORF prediction and evidence for translation activity from ribo-seq data analysis

We first annotated all ORFs across entire genomes using RibORF (Ji 2018). A minimal length of nine nucleotides was specified, and non-canonical START codons were allowed: TTG, CTG, GTG and ACG. We classified ORFs as IGORFs if they display no overlap with any protein-coding gene annotation.

To obtain evidence for translation activity across the genome (and IGORFs in particular), we then analysed the ribo-seq data. Adapters were trimmed from the reads and then filtered on quality and length using Cutadapt (v3.2) (Martin 2011). We removed known non-coding RNAs such as rRNA, tRNA and snoRNA through bowtie2 (v2.4.1, Langmead and Salzberg 2012) alignments on TAIR10 non-coding RNA annotations. Depleted sequences were then mapped on the corresponding reference genome per species: AUBY1 for *A. halleri* (Pavan et al. 2025) and MN47 for *A. lyrata* (Kolesnikova et al. 2023). The alignment was performed using the splice-aware aligner STAR (v2.7.5a) (Dobin et al. 2013), tolerating up to one mismatch in the alignment, and up to 20 multi-mapping positions. For *A. thaliana*, raw sequencing data from (Hsu et al. 2016; Wu and Hsu 2022) (GEO accession GSE81332 and BioProject ID PRJNA854638) were trimmed from their adapters, filtered on quality and length using Cutadapt (same parameters as previously) and further processed identically to the data produced in this study, using the TAIR10 reference genome (Lamesch et al. 2012). The quality of the ribosome profiling datasets was assessed by producing protein-coding gene meta-plots using RibORF (Ji 2018) to visualize the periodicity signals. We inferred the P-site offsets by visual assessment of the metagene plots. The mapped reads were subsequently shifted by the assigned offsets using RibORF.

Signatures of translation for each annotated CDS and IGORFs within those candidates were detected by three methods. First we used ORFSCORE (Bazzini et al. 2014), where ribosome footprints accumulation is computed in each possible frame, with the assumption that coding nucleotide sequences would exhibit greater accumulation in the first frame. Annotations were produced using the R (v4.2.2) packages ORFik (Tjeldnes et al. 2021), rtracklayer (Lawrence et al. 2009), GenomicRanges and GenomicFeatures (Lawrence et al. 2013). Predicted ORFs that fell within the 95% confidence interval of the CDS ORFscore distribution were retained. Second, we used RibORF, which relies on Machine Learning (“Support Vector Machine”), trained to recognize triplet periodicity on canonical gene annotations specific to each species. Predictions were performed on candidates with a cut-off that was adjusted to each dataset to keep a FDR < 0.05. Finally, we used RiboCode (Xiao et al. 2018), which allows for the *de novo* annotation of periodicity signals using a modified Wilcoxon signed-rank test. The default p-value (0.001) was used to distinguish translated sequences.

### Syntenic region annotation and alignment

CDS annotations for each species were used to identify gene orthologs between all pairs of species using Inparanoid (Remm et al. 2001), and keeping pairs of orthologs with a score ≥ 100 and a bootstrap value ≥ 99. Syntenic regions were consequently annotated with homemade R scripts, where a syntenic region was defined as a genomic region anchored by two conserved CDS orthologs, and which includes intergenic sequences. Non-conserved CDSs and IGORFs that fell within syntenic regions were identified with the bedtools suite “intersect” (Quinlan and Hall 2010). An initial local pairwise alignment was performed between each non-conserved CDS, or IGORF, and its corresponding syntenic region using EMBOSS water (Madeira et al. 2024), and used to identify the coordinates of the orthologous region in the pair species. Coordinates of the non-conserved CDS, or IGORF region were then extended 1kb up- and downstream (or until the closest conserved CDS annotation), and were globally aligned using the ggsearch36 tool from the fasta36 software (v3.8i) (Pearson and Lipman 1988) with an e-value threshold of 0.01. In parallel, the non-conserved CDSs and IGORFs falling within syntenic regions were used for a BLASTX search (Camacho et al. 2009) against the proteomes of five *Brassicaceae* species downloaded from Genbank (Benson et al. 2012): *Camelina sativa*, *Brassica napus*, *Brassica oleraceae*, *Eutrema salsugineum* and *Raphanus sativus*. CDSs and IGORFs that showcased any significant hit were discarded (threshold e- value of 0.001). The CDSs and IGORFs that were kept were subjected to a BLASTn search (Camacho et al. 2009) against genomes of every species of interest. The ones that exhibited a significant hit (default e-value) outside of the expected syntenic region, and whose length was ≥ 50% of the CDS or IGORF’s length, were discarded. Among the remaining sequences, the Percent Identity (ID) between each non-conserved CDS, or IGORF, and all its syntenic aligned regions were computed. Two elements were required to select CDSs and IGORFs with sufficient homology: that the global pairwise alignment was significant with *C. rubella*, or if not with *A. thaliana*, and that the PID in the CDS/IGORF region within the pairwise alignment was ≥ 0.6. CDSs and IGORFs that passed those criteria were used for Multiple Sequence Alignment (MSA); between the extended CDS/IGORF sequence and every correctly aligned, extended syntenic region using MAFFT (Katoh 2002). Once the multiple alignments were obtained, they were used for conservation analyzes with the help of home-made python scripts. Multiple alignments were further filtered, with a PID of at least 0.5 in at least three species, and a maximal gap content of 40%, all within the CDS/IGORF region. Finally, a minimal length of 30 nucleotides was required and smaller CDS/IGORFs were discarded. The *de novo* status was inferred when a region lacking CDS/IGORF conservation (START, STOP and reading frame) could be identified, or if a CDS/IGORF had not been bioinformatically predicted at those positions, in either *C. rubella* or in *A. thaliana*. Furthermore, if in any pair species a CDS/IGORF was absent but its position overlapped with a gene annotation, the conservation of the frame was checked, and the sequence was considered as conserved if the gene’s frame was the same as that of the reference CDS/IGORF.

### Intrinsic disorder and structure prediction

Alphafold2 (v.2.3.0) (Jumper et al. 2021) predictions were performed on HPC cluster Palma II. Predictions with highest confidence (ranked_0) were used for analysis. Disorder was predicted locally using AIUpred (no smoothing) (Erdős and Dosztányi 2024). Mean and median disorder per sequence were calculated using Python v3.10.13 with pandas v2.1.1 (McKinney 2011). Structures were visualized using UCSF ChimeraX-1.10.1 (Pettersen et al. 2021).

### Cross-accession conservation

We gathered five high-quality *A. lyrata* genome assemblies (PTP, JBTAFC000000000), INDP15 and TSS14.18 (ENA project: PRJEB80457 (Padilla-García et al. 2025)), GCA_026151145 (GenBank assembly accession: GCA_026151145.1) from Austria, and NT1 (GenBank assembly accession: GCA_963992905.1 from Siberia (Scott et al. 2025)). Annotations from the reference genomes of *A. lyrata* (MN47) were lifted off to the other assemblies using the liftoff tool (v1.6.3) (Shumate and Salzberg 2021). Using the same parameters as previously, all potential CDSs and IGORFs candidates were bioinformatically annotated with RibORF (Ji 2018), on the basis of their sequence only, in each accession. The coordinates of the syntenic region from the reference genome were transposed to each accession, and conserved syntenic regions in different accessions were aligned using MAFFT (Katoh 2002). The conservation of the CDS or IGORF sequence was then assessed from these multiple alignments, using the same procedure as previously. To test whether translated *de novo* IGORFs/CDSs or untranslated intergenic ORFs were more conserved across accessions, we created a “conservation score” that is, per CDS or IGORF, the number of accessions within which each IGORF/CDS is found. A Mann-Whitney U test was performed across the two categories for the observed conservation patterns, and “*de novo”* and “untranslated IGORF” labels were then randomly permuted 1,000 times, with new conservation scores and a new Mann-Whitney U test being computed for each iteration. The observed test score was then compared to the distribution of test scores under the null hypothesis (*de novo* and untranslated IGORFs have similar conservation pattern), and a corresponding p-value was assessed.

### Polymorphism analysis

Genomic data from a total of 39 *A. lyrata* individuals from North America and central Europe (**Table S7**) (Mattila et al. 2017; Hämälä et al. 2018; Marburger et al. 2019; Takou et al. 2021; Pavan et al. 2025) were collected. Paired-end reads were mapped to the MN47 *A. lyrata* reference genome using bowtie 2 (v2.4.1) (Langmead and Salzberg 2012) and duplicates were marked and removed using Picard Tools (v2.21.4) (Anon 2019). Variant calling and subsequent genotype joining were performed using GATK (v4.1.9.0) (Van der Auwera and O’Connor 2020). Variant calling files were split into invariant and filtered variant sites (filtered with VCFtools (Danecek et al. 2011), with parameters *--mac 1, --remove-indels, --minDP 10, --min-alleles 1, --max-alleles 2, --max-missing 0.75* and *--minQ 30*), and concatenated back into one single variant calling file with BCFtools (Danecek et al. 2021). Average nucleotide diversity per site was computed from the concatenated variant calling file, within each feature type, using pixy (Korunes and Samuk 2021). Bedtools intersect (Quinlan and Hall 2010) was used to retrieve regions of the variant calling file corresponding to each individual annotation, which were then converted into fasta sequences for each allele of each individual using BCFtools consensus (Danecek et al. 2021) with default parameters. The resulting sequences were aligned using MAFFT (Katoh 2002). Because of the limited number of polymorphic sites in *de novo* ORFs and unt_IGORFs resulting from their short lengths, all alignments from each category were concatenated. To get an equivalent number of polymorphic sites, five randomly selected conserved CDS alignments were concatenated and the procedure was repeated 1,000 times. The *π*N/*π*S ratios were computed from these concatenated alignments using dNdSpiNpiS_1.0 (Gayral et al. 2013) with parameters *-gapN_site=10*, *-gapN_seq=0.6*, *- remove_frameshift=yes* and *-allow_internal_bc=yes*; and the *π*N/*π*S ratios of conserved CDS from the 1,000 samples were plotted as a distribution.

Plots were produced using R (v 4.2.2) (R Core Team 2025), with packages ggplot2 (Wickham 2009), ggsignif (Constantin and Patil 2021), ggpubr (Kassambara 2023), patchwork (Pedersen 2024), ComplexUpset (Michał Krassowski et al. 2022) and ComplexHeatmap (Gu et al. 2016), and alignment visualizations with Jalview (Waterhouse et al. 2009). *De novo* ORF annotations are available on Zenodo at https://zenodo.org/records/18230755, as well as the DeNoFo standardized annotation (Dohmen et al. 2025).

## Acknowledgements

We acknowledge the sequencing and bioinformatics expertise of the I2BC High-throughput sequencing facility, supported by France Génomique (funded by the French National Program “Investissement d’Avenir” ANR-10-INBS-09), as well as Anne Lopes for sharing with us her expertise in Ribosome Profiling data analysis. We thank the high performance computing service and Bilille at the University of Lille for providing computing resources, as well as Mathieu Genete and Clément Mazoyer for providing bioinformatic support. This work was performed using the infrastructure and technical support of the “Plateforme Serre, cultures et terrains expérimentaux – Université de Lille” for the greenhouse/field facilities, and we thank Pierre Saumitou and Flavia Pavan for their help in setting up the hydroponic culture protocol. Structure predictions for this publication were performed on the HPC cluster PALMA II of the University of Münster, subsidized by the Deutsche Forschungsgemeinschaft (INST 211/667-1).

## Data availability

Scripts used for the analyses presented in the paper are available at https://zenodo.org/records/21535083

Ribosome profiling sequencing reads from *A. lyrata* and *A. halleri* are available under the BioProject PRJNA1473080.

