## Supplementary materials for "Pervasive translation of intergenic open reading frames and *de novo* gene emergence in *Arabidopsis*"


**Authors**

Claire Patiou[1], Christelle Blassiau[1], Isabelle Hatin[2], Lars A. Eicholt[3], Enora Corler[2], Chloé Ponitzki[1], Laurence Debacker[1], Erich Bornberg-Bauer[3,4], Olivier Namy[2], Sylvain Legrand[1], Vincent Castric[1], Eléonore Durand[1]

[1] Univ. Lille, CNRS, UMR 8198 - Evo-Eco-Paleo, F-59000 Lille, France

[2]Université Paris-Saclay, CEA, CNRS, Institute for Integrative Biology of the Cell (I2BC), 91198 Gif-sur-Yvette, France

[3] Institute for Evolution and Biodiversity, University of Muenster, 48149 Muenster, Germany

[4]Department of Protein Evolution, Max-Planck Institute for Biology, Max-Planck-Ring 5, 72076 Tuebingen, Germany

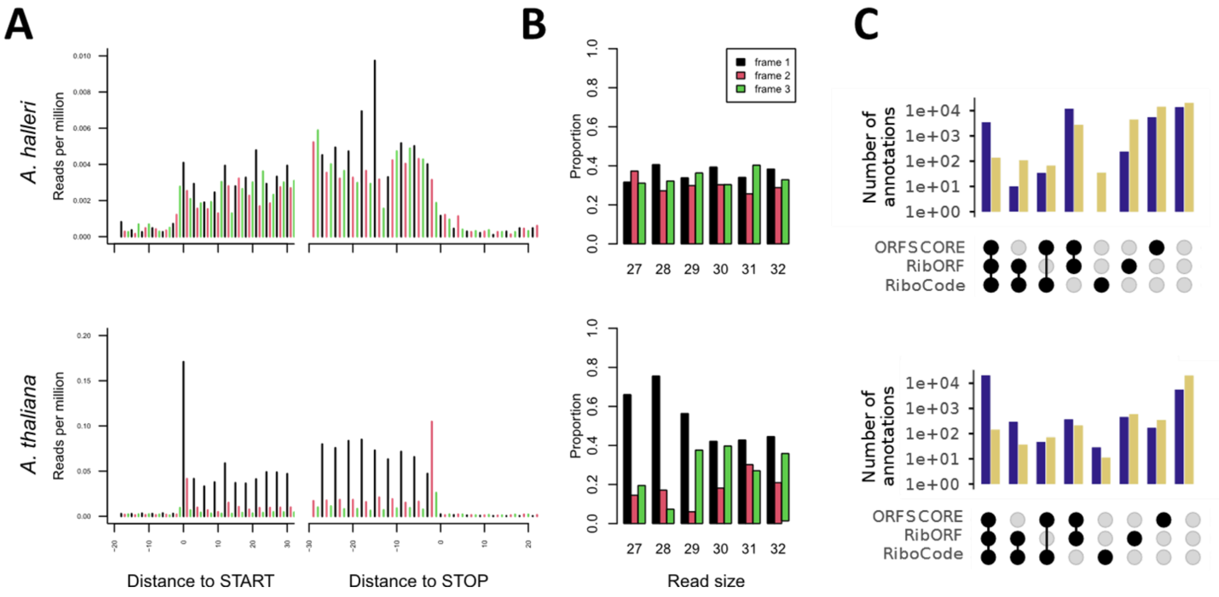


**Figure S1. Detection of translated CDSs and intergenic ORFs (IGORFs) in *A. halleri* (top) and *A. thaliana* (bottom). A)** Metagene plots of 28-kmers from ribosome profiling, after P-site shifting (reads were shifted according to their k-mer specific P-site offset, and trimmed to be 1 base-long), used for the annotation of translation events; and prevalence of k-mers depending on which frame they are found in, produced with modified RibORF scripts (Ji 2018). Metagene plots are created by concatenating the accumulation of ribosome profiling footprints within all known canonical protein-coding genes, measured in reads per million (RPM) relative to the start position of CDS. Read accumulation is colored based on its frame relative to the CDS: black for the first frame, red for the second frame and green for the third frame. **B)** The relative abundance of each k-mer, between 27 and 32 nucleotides in size, is also colored based on the frame in which the reads are located. Ribosomal footprints are expected to be around 28 nucleotides long, i.e. the length of the mRNA segment protected by the ribosome (Ingolia et al. 2009). **C)** Number of annotations detected as translated ($log_{10}$ scale) according to the tools that detected them as translated, and to the category to which they belong: canonical CDS or IGORFs. Black circles represent the detection of translation, grey circles represent its absence.

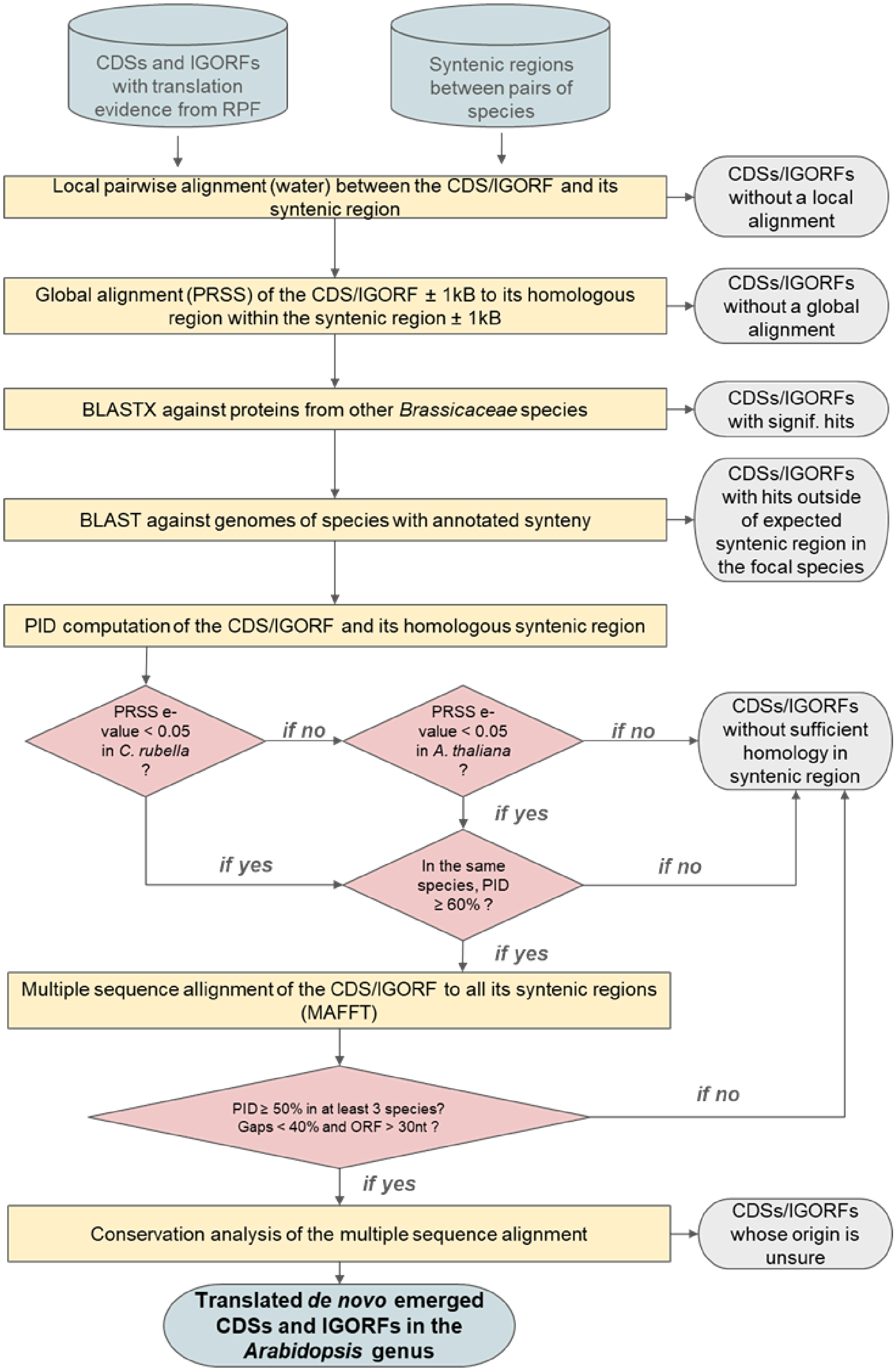


**Figure S2:** Flowchart of the bioinformatic pipeline used for the annotation of translated de novo CDSs and IGORFs.

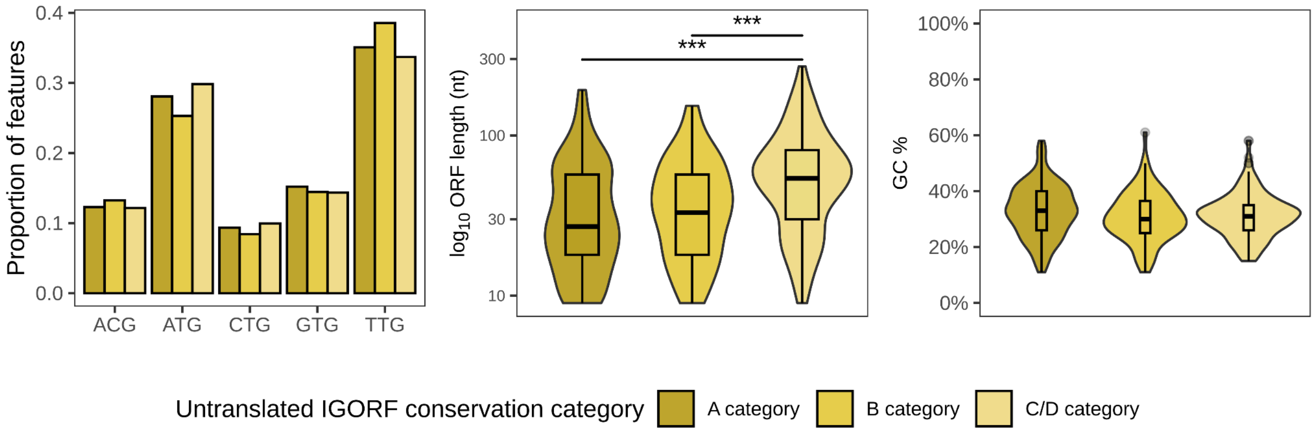


**Figure S3:** Sequences properties of untranslated IGORFs (START codon, length and GC content) depending on their conservation class. For length and GC content, all statistically significant differences were represented (Mann-Whitney U-test, * = p < 0.05, ** = p < 0.01, *** = p< 0.001).

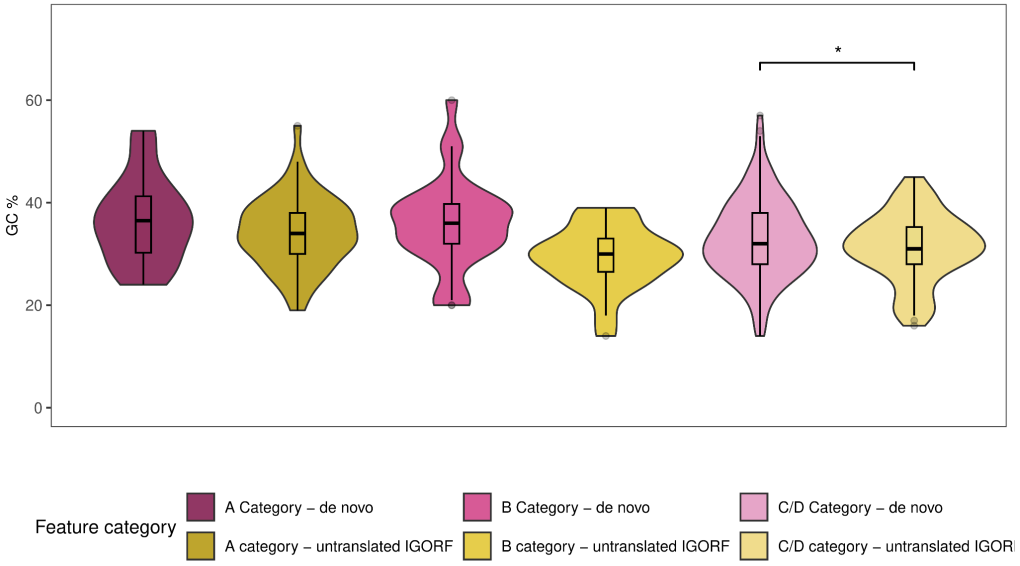


**Figure S4:** GC content of untranslated IGORs and *de novo* ORFs depending on their conservation class. All statistically significant differences were represented (Mann-Whitney U-test, * = p < 0.05, ** = p < 0.01, *** = p< 0.001).

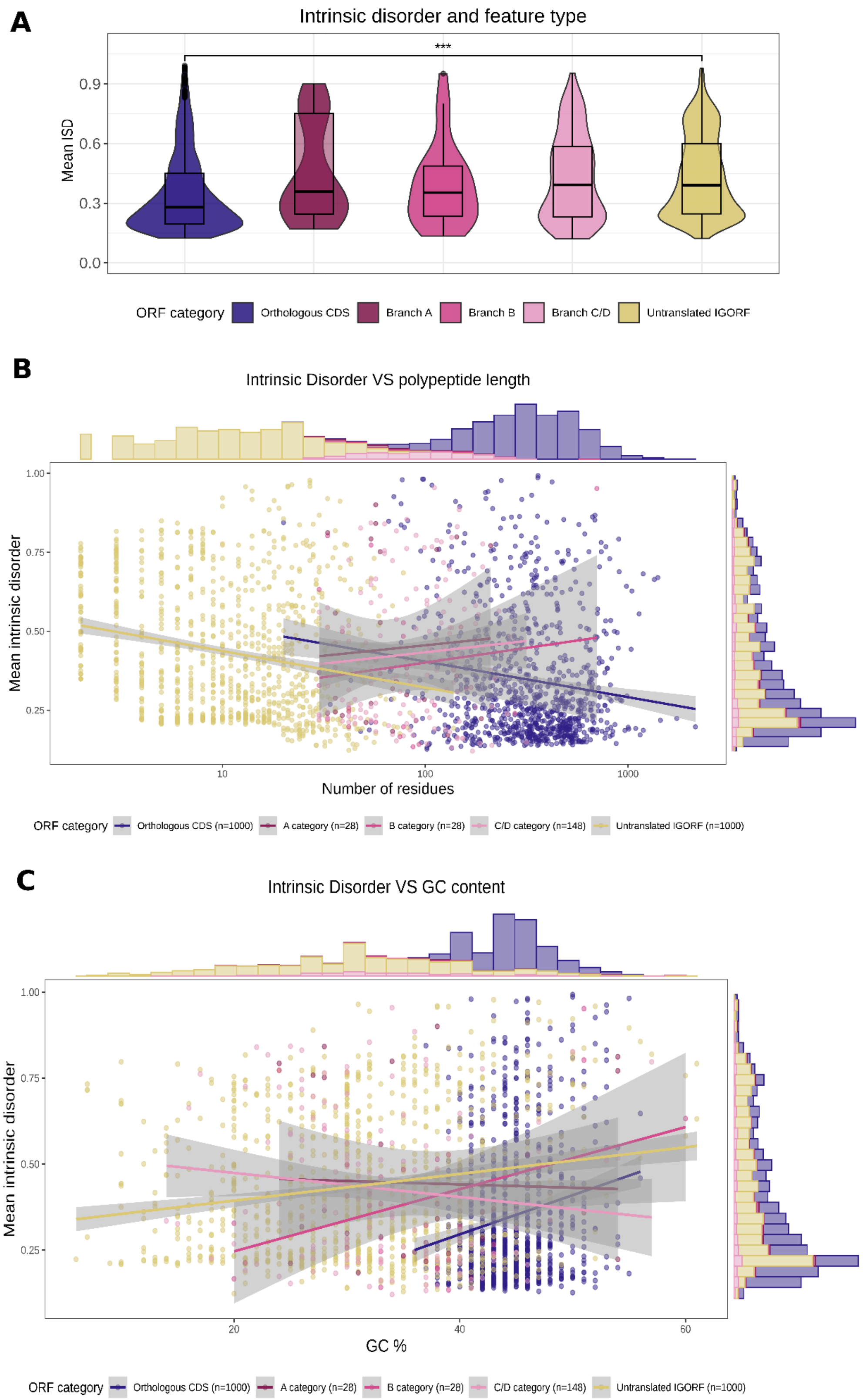
A
Intrinsic disorder and feature type
***
Mean ISD
0.0
0.3
0.6
0.9
ORF category
Orthologous CDS
Branch A
Branch B
Branch C/D
Untranslated IGORF
B
Intrinsic Disorder VS polypeptide length
Mean intrinsic disorder
0.25
0.50
0.75
1.00
10
100
1000
Number of residues
ORF category
Orthologous CDS (n=1000)
A category (n=28)
B category (n=28)
C/D category (n=148)
Untranslated IGORF (n=1000)
C
Intrinsic Disorder VS GC content
Mean intrinsic disorder
0.25
0.50
0.75
1.00
20
40
60
GC %
ORF category
Orthologous CDS (n=1000)
A category (n=28)
B category (n=28)
C/D category (n=148)
Untranslated IGORF (n=1000)

**Figure S5**: **A)** Measures of ISD in the translation products of conserved canonical CDS, different conservation classes of *de novo* ORFs and untranslated IGORFs. The only statistically significant difference (between conserved CDS and untranslated IGORFs) is represented (Mann-Whitney U-test, *** = p< 0.001). **B)** Linear regression of Mean Intrinsic Disorder in the translation products of the same previous categories, against the number of amino acid residues. Across all categories a significant negative association was found between the 2 variables (*adjusted* $R^2 = 0.03532$, *p-value* $< 2.2\times 10^{-16}$), as well as within unt_IGORFs and conserved CDS (*unt_IGORFs: Spearman* $\rho$*(disorder,length)= -0.24, p-value=* $1.22\times 10^{-14}$*; conserved CDS:* $\rho$*(disorder,length)= -0.14, p-value=*$1.29\times 10^{-5}$). No significant association was found between these 2 variables within and across *de novo* ORF conservation categories. **C)** Linear regression of Mean Intrinsic Disorder in the translation products of the same previous categories against the level of GC content in the DNA sequences encoding them. A significant positive association was found across annotations (*adjusted* $R^2 = 0.0844$, $p < 2.2 \times 10^{-16}$), as well as within unt_IGORFs and conserved CDS (*unt_IGORFs: Spearman* $\rho$*(disorder,GC)= 0.13, p-value=* $2.53\times 10^{-5}$*; conserved CDS:* $\rho$*(disorder,GC)= 0.15, p-value=*$1.67\times 10^{-6}$). A negative correlation between ISD and GC was found in C/D category *de novo* ORFs ($\rho$*(disorder,GC)= -0.20, p-value=0.01474*).

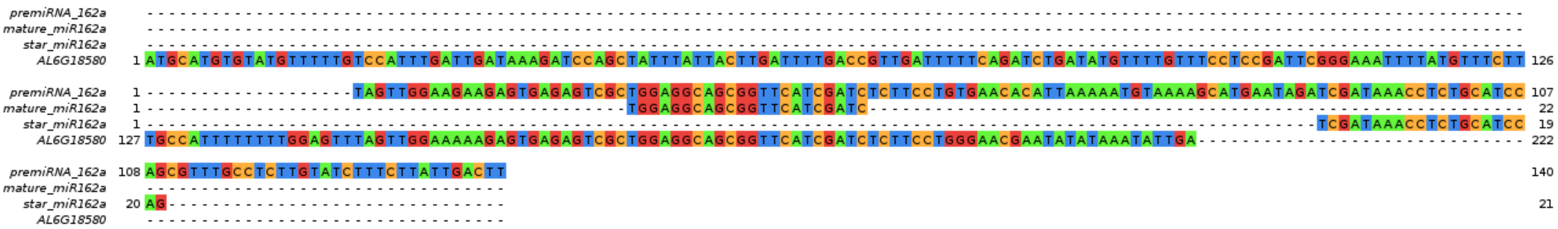


**Figure S6:** Alignment of the *A. lyrata* AL6G18580 gene to the *A. thaliana* miR162a precursor, mature miRNA and miRNA* (star miRNA).

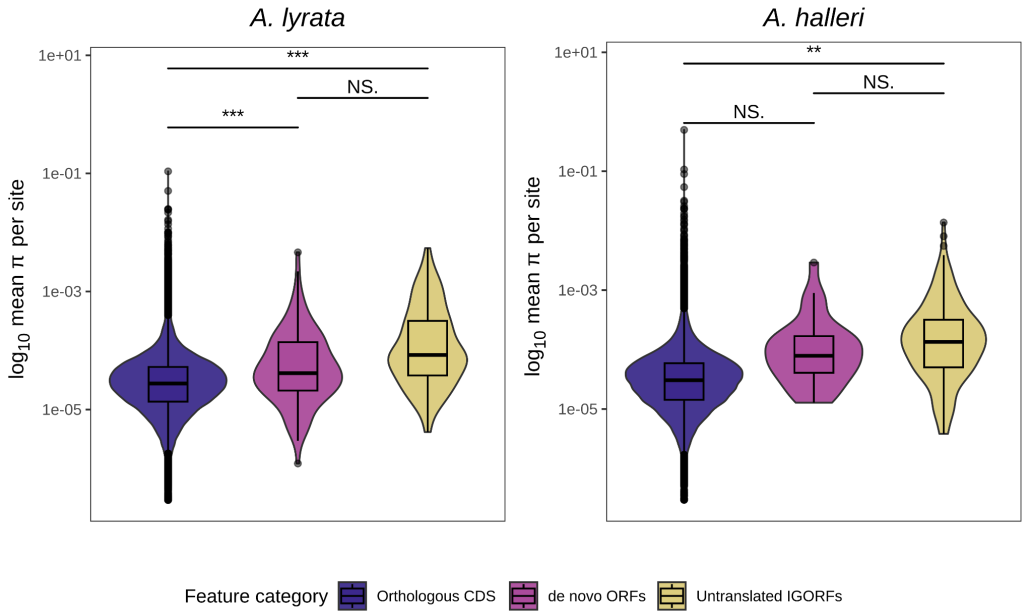


**Figure S7:** Average nucleotide diversity per site, per feature category: conserved CDS, translated *de novo* ORs or untranslated IGORFs (Mann-Whitney U-test, * = $p < 0.05$, ** = $p < 0.01$, *** = $p < 0.001$).

| | Annotation number | | Detection of translation by: | | | Translated annotation number | |
|---|---|---|---|---|---|---|---|
| Species | Canonical genes | IGORF | RiboCode | RibORF | ORFSCORE | Canonical genes | IGORF |
| *A. lyrata* | 30 443 | 23 503 084 | YES | YES | YES | 13 266 | 1 548 |
| | | | NO | YES | YES | 8 210 | 7 114 |
| | | | YES | NO | YES | 101 | 676 |
| | | | YES | YES | NO | 9 | 292 |
| | | | NO | NO | YES | 2 701 | 60 249 |
| | | | NO | YES | NO | 108 | 15 684 |
| | | | YES | NO | NO | 6 | 1 870 |
| | | | | | **TOTAL:** | **24 401** | **87 433** |
| *A. halleri* | 34 805 | 19 117 874 | YES | YES | YES | 3 467 | 135 |
| | | | NO | YES | YES | 11 952 | 2 727 |
| | | | YES | NO | YES | 34 | 66 |
| | | | YES | YES | NO | 10 | 106 |
| | | | NO | NO | YES | 5 466 | 14 195 |
| | | | NO | YES | NO | 238 | 4 424 |
| | | | YES | NO | NO | 0 | 35 |
| | | | | | **TOTAL:** | **21 167** | **21 688** |
| *A. thaliana* | 27 559 | 3 327 672 | YES | YES | YES | 20 592 | 142 |
| | | | NO | YES | YES | 364 | 208 |
| | | | YES | NO | YES | 47 | 70 |
| | | | YES | YES | NO | 295 | 36 |
| | | | NO | NO | YES | 169 | 346 |
| | | | NO | YES | NO | 451 | 583 |
| | | | YES | NO | NO | 28 | 11 |
| | | | | | **TOTAL:** | **21 946** | **1 396** |

**Table S1:** Summary of the data used to produce Figures 1 and S1: the number of available annotations, the number of tools detecting them as translated, and the resulting number of annotations found as translated.

| | Sample number | Raw reads | Reads after adapter trimming | Reads after quality control and size selection | Reads after *in silico* small rRNA depletion | % of removed residual rRNAs | Reads mapped once to genome* | Reads mapped 2-20 times to genome* | Mapped too many times (>20) | Unmapped | Reference |
|---|---|---|---|---|---|---|---|---|---|---|---|
| *A. lyrata* | ALY | 609 771 764 | 601 833 807 | 409 856 100 | 242 989 105 | 41 | 22 947 667 | 8 584 665 | 5 704 766 | 195 785 778 | This study |
| *A. halleri* | AHA | 639 109 284 | 614 402 572 | 168 360 788 | 114 000 850 | 32 | 3 409 600 | 14 812 250 | 36 770 015 | 58 882 173 | This study |
| *A. thaliana* | Root_1 | 163 396 264 | 159 778 193 | 142 176 011 | 107 573 216 | 24 | 73 679 997 | 19 213 813 | 18 778 | 14 625 936 | Hsu *et al*., 2016 |
| | Root_2 | 166 148 265 | 162 112 008 | 138 493 159 | 115 260 206 | 17 | 71 582 384 | 13 005 304 | 18 222 | 30 625 465 | Hsu *et al*., 2016 |
| | Root_3 | 207 346 547 | 202 257 844 | 137 022 278 | 97 879 433 | 29 | 69 194 868 | 19 289 223 | 39 751 | 9 299 834 | Hsu *et al*., 2016 |
| | Shoot_1 | 145 235 719 | 142 483 567 | 121 767 293 | 102 210 257 | 16 | 33 727 185 | 11 065 144 | 20 256 | 57 388 016 | Hsu *et al*., 2016 |
| | Shoot_2 | 179 566 257 | 175 969 706 | 163 389 872 | 129 057 937 | 21 | 87 963 741 | 22 356 181 | 23 358 | 18 682 755 | Hsu *et al*., 2016 |
| | Shoot_3 | 134 241 510 | 131 386 200 | 117 584 678 | 84 372 098 | 28 | 59 378 977 | 11 417 693 | 20 030 | 13 535 155 | Hsu *et al*., 2016 |
| | Seedlings | 185 867 719 | 183 322 013 | 87 331 306 | 66 719 312 | 24 | 32 452 595 | 32 075 787 | 44 822 | 2 141 266 | Hsu *et al*., 2022 |

**Table S2:** Summary of the ribosome profiling sequencing data, both from this study and from publicly available datasets. Columns marked with a * correspond to reads used to annotate translation events.

| | Species pair | Translated ORFs found within syntenic regions | Without significant hits in protein coding genes of other *Brassicaceae* species | Without significant hits in genomic regions outside of synteny in focal species | With an extended alignment in *C. rubella* or *A. thaliana* of E-value < 0.05 and a PID > 60% | With a MSA of at least 50% PID in at least 3 sequences, and maximum of 40% gaps | With a *de novo* emergence |
|---|---|---|---|---|---|---|---|
| *A. halleri* | *A. lyrata* | 2,880 | 1,416 | 864 | 302 | 192 | 42 |
| | *A. thaliana* | 3,334 | | | | | |
| | *C. rubella* | 3,100 | | | | | |
| *A. lyrata* | *A. halleri* | 7,152 | 5,454 | 3,520 | 1,282 | 697 | 163 |
| | *A. thaliana* | 8,614 | | | | | |
| | *C. rubella* | 8,504 | | | | | |
| *A. thaliana* | *A. halleri* | 2,192 | 376 | 247 | 39 | 31 | 14 |
| | *A. lyrata* | 2,362 | | | | | |
| | *C. rubella* | 2,888 | | | | | |

**Table S3:** Summary of the data produced by synteny analyzes as shown in Figure S3.

| ORF name | Hits description | Query Cover | E value | Per. ident |
|---|---|---|---|---|
| GENE_732871 | A. lyrata uncharacterized LOC110227695 mRNA,<br>A. thaliana uncharacterized protein AT5G12043,<br>A. thaliana genome | 99%,<br>100%,<br>100% | 3E-33,<br>2E-25,<br>2E-25 | 87.94,<br>84.51,<br>84.51 |
| GENE_826985 | A. lyrata probable apyrase 7 LOC9304022 mRNA,<br>A. thaliana genome,<br>A. thaliana GDA1/CD39 nucleoside phosphatase family protein AT4G19180 mRNA | 100%,<br>62%,<br>62% | 7E-40,<br>6.00E-31,<br>3E-29 | 87.8,<br>95.6,<br>94.51 |
| GENE_829626 | NO_HITS | NO_HITS | NO_HITS | NO_HITS |
| GENE_AL1G18050 | NO_HITS | NO_HITS | NO_HITS | NO_HITS |
| GENE_AL3G13390 | NO_HITS | NO_HITS | NO_HITS | NO_HITS |
| GENE_AL3G23310 | A. lyrata uncharacterized LOC9318812 mRNA,<br>A. thaliana genome<br>A. thaliana uncharacterized protein AT3G11745 mRNA | 100%,<br>100%,<br>100% | 0.0000006,<br>4E-26,<br>4E-26 | 100,<br>88.68,<br>88.68 |
| GENE_AL3G25260 | A. thaliana genome,<br>A. thaliana uncharacterized misc_RNA (AT3G00320) | 73%,<br>73% | 0.00000007,<br>0.00000007 | 100,<br>100 |
| GENE_AL4G44240 | A. lyrata uncharacterized LOC9318064 mRNA | 100% | 2.00E-49 | 100 |
| GENE_AL4G45690 | A. thaliana genome | 52% | 4.00E-17 | 92.86 |
| GENE_AL5G37070 | A. lyrata glycine-rich protein 3 short isoform LOC9312403 mRNA | 100% | 5.00E-92 | 100 |
| GENE_AL6G11140 | A. thaliana genome,<br>A. thaliana uncharacterized misc_RNA AT3G00320 | 86%,<br>86% | 5E-30,<br>5E-30 | 92.86,<br>92.86 |
| GENE_AL6G18580 | A. lyrata uncharacterized LOC110227896 ncRNA,<br>A. lyrata microRNA aly-MIR162a precursor,<br>A. thaliana miR162a primary transcript gene,<br>Arabidopsis thaliana hypothetical protein MIR162a miRNA | 100%,<br>54%,<br>78%,<br>27% | 3E-110,<br>1E-53,<br>2E-42,<br>7E-17 | 100,<br>86.26,<br>86.26,<br>96.67 |
| GENE_AL8G35520 | A. lyrata extensin-3 LOC9302389 mRNA,<br>A. thaliana Proline-rich extensin-like family protein AT5G59170 mRNA,<br>A. thaliana genome | 97%,<br>88%,<br>88% | 0,<br>8E-86,<br>6E-14 | 100,<br>87.76,<br>96.23 |
| IGORF_1 | A. thaliana genome | 100% | 2.00E-60 | 100 |
| IGORF_2 | A. thaliana genome | 100% | 3.00E-95 | 100 |
| IGORF_3 | NO_HITS | NO_HITS | NO_HITS | NO_HITS |
| IGORF_4 | A. thaliana genome | 100% | 6.00E-26 | 100 |
| IGORF_5 | NO_HITS | NO_HITS | NO_HITS | NO_HITS |
| IGORF_6 | A. thaliana genome,<br>B. rapa genome | 100%,<br>42% | 6E-44,<br>3.00E-07 | 100,<br>95.35 |
| IGORF_7 | A. thaliana genome | 100% | 2.00E-32 | 100 |
| IGORF_8 | A. thaliana genome | 100% | 1.00E-103 | 100 |
| IGORF_9 | A. thaliana genome | 100% | 2.00E-32 | 100 |
| IGORF_10 | A. thaliana genome | 100% | 2.00E-19 | 100 |
| IGORF_12 | A. thaliana genome,<br>A. lyrata uncharacterized LOC9310279 misc_RNA | 100%,<br>100% | 2E-42,<br>3.00E-17 | 100,<br>86 |
| IGORF_14 | A. thaliana genome | 100% | 3.00E-16 | 100 |
| IGORF_15 | NO_HITS | NO_HITS | NO_HITS | NO_HITS |
| IGORF_16 | NO_HITS | NO_HITS | NO_HITS | NO_HITS |
| IGORF_17 | A. thaliana genome | 98% | 7.00E-56 | 89.84 |
| IGORF_18 | NO_HITS | NO_HITS | NO_HITS | NO_HITS |
| IGORF_19 | NO_HITS | NO_HITS | NO_HITS | NO_HITS |
| IGORF_20 | NO_HITS | NO_HITS | NO_HITS | NO_HITS |
| IGORF_22 | NO_HITS | NO_HITS | NO_HITS | NO_HITS |
| IGORF_23 | NO_HITS | NO_HITS | NO_HITS | NO_HITS |
| IGORF_26 | A. thaliana genome | 100% | 2.00E-14 | 96.3 |
| IGORF_27 | NO_HITS | NO_HITS | NO_HITS | NO_HITS |
| IGORF_28 | NO_HITS | NO_HITS | NO_HITS | NO_HITS |
| IGORF_29 | NO_HITS | NO_HITS | NO_HITS | NO_HITS |
| IGORF_30 | NO_HITS | NO_HITS | NO_HITS | NO_HITS |
| IGORF_32 | NO_HITS | NO_HITS | NO_HITS | NO_HITS |
| IGORF_33 | NO_HITS | NO_HITS | NO_HITS | NO_HITS |
| IGORF_34 | NO_HITS | NO_HITS | NO_HITS | NO_HITS |
| IGORF_35 | NO_HITS | NO_HITS | NO_HITS | NO_HITS |
| IGORF_36 | NO_HITS | NO_HITS | NO_HITS | NO_HITS |
| IGORF_37 | NO_HITS | NO_HITS | NO_HITS | NO_HITS |
| IGORF_38 | NO_HITS | NO_HITS | NO_HITS | NO_HITS |
| IGORF_40 | NO_HITS | NO_HITS | NO_HITS | NO_HITS |
| IGORF_42 | NO_HITS | NO_HITS | NO_HITS | NO_HITS |
| IGORF_43 | NO_HITS | NO_HITS | NO_HITS | NO_HITS |
| IGORF_44 | NO_HITS | NO_HITS | NO_HITS | NO_HITS |
| IGORF_45 | NO_HITS | NO_HITS | NO_HITS | NO_HITS |
| IGORF_46 | NO_HITS | NO_HITS | NO_HITS | NO_HITS |
| IGORF_47 | NO_HITS | NO_HITS | NO_HITS | NO_HITS |
| IGORF_48 | Sisymbrium irio genome | 30% | 0.0002 | 100 |

| | | | | |
|---|---|---|---|---|
| IGORF_49 | NO_HITS | NO_HITS | NO_HITS | NO_HITS |
| IGORF_50 | NO_HITS | NO_HITS | NO_HITS | NO_HITS |
| IGORF_52 | NO_HITS | NO_HITS | NO_HITS | NO_HITS |
| IGORF_53 | NO_HITS | NO_HITS | NO_HITS | NO_HITS |
| IGORF_55 | NO_HITS | NO_HITS | NO_HITS | NO_HITS |
| IGORF_56 | NO_HITS | NO_HITS | NO_HITS | NO_HITS |
| IGORF_57 | NO_HITS | NO_HITS | NO_HITS | NO_HITS |
| IGORF_58 | Brassica oleracea genome | 30% | 3.00E-08 | 97.56 |
| IGORF_61 | NO_HITS | NO_HITS | NO_HITS | NO_HITS |
| IGORF_63 | A. thaliana genome | 100% | 3.00E-25 | 95.06 |
| IGORF_65 | A. lyrata uncharacterized LOC9325856 mRNA | 100% | 2.00E-08 | 100 |
| IGORF_66 | NO_HITS | NO_HITS | NO_HITS | NO_HITS |
| IGORF_67 | NO_HITS | NO_HITS | NO_HITS | NO_HITS |
| IGORF_68 | NO_HITS | NO_HITS | NO_HITS | NO_HITS |
| IGORF_69 | NO_HITS | NO_HITS | NO_HITS | NO_HITS |
| IGORF_70 | NO_HITS | NO_HITS | NO_HITS | NO_HITS |
| IGORF_71 | NO_HITS | NO_HITS | NO_HITS | NO_HITS |
| IGORF_72 | NO_HITS | NO_HITS | NO_HITS | NO_HITS |
| IGORF_73 | NO_HITS | NO_HITS | NO_HITS | NO_HITS |
| IGORF_74 | NO_HITS | NO_HITS | NO_HITS | NO_HITS |
| IGORF_75 | A. thaliana genome | 86% | 3.00E-18 | 84.96 |
| IGORF_76 | NO_HITS | NO_HITS | NO_HITS | NO_HITS |
| IGORF_77 | NO_HITS | NO_HITS | NO_HITS | NO_HITS |
| IGORF_78 | NO_HITS | NO_HITS | NO_HITS | NO_HITS |
| IGORF_79 | A. thaliana genome | 65% | 1.00E-14 | 80 |
| IGORF_80 | NO_HITS | NO_HITS | NO_HITS | NO_HITS |
| IGORF_81 | NO_HITS | NO_HITS | NO_HITS | NO_HITS |
| IGORF_82 | NO_HITS | NO_HITS | NO_HITS | NO_HITS |
| IGORF_83 | NO_HITS | NO_HITS | NO_HITS | NO_HITS |
| IGORF_84 | Brassica oleracea genome | 88% | 1.00E-08 | 100 |
| IGORF_85 | NO_HITS | NO_HITS | NO_HITS | NO_HITS |
| IGORF_86 | NO_HITS | NO_HITS | NO_HITS | NO_HITS |
| IGORF_87 | A. thaliana genome | 100% | 5.00E-34 | 96.81 |
| IGORF_88 | A. thaliana genome,<br>A. thaliana other RNA AT1G51645 lncRNA | 96%,<br>96% | 1E-48,<br>1E-48 | 87.36<br>87.36 |
| IGORF_89 | NO_HITS | NO_HITS | NO_HITS | NO_HITS |
| IGORF_90 | A. thaliana genome | 72% | 1.00E-27 | 91.84 |
| IGORF_92 | NO_HITS | NO_HITS | NO_HITS | NO_HITS |
| IGORF_93 | A. thaliana genome | 51% | 1.00E-39 | 95.5 |
| IGORF_94 | NO_HITS | NO_HITS | NO_HITS | NO_HITS |
| IGORF_96 | NO_HITS | NO_HITS | NO_HITS | NO_HITS |
| IGORF_97 | NO_HITS | NO_HITS | NO_HITS | NO_HITS |
| IGORF_98 | NO_HITS | NO_HITS | NO_HITS | NO_HITS |
| IGORF_100 | NO_HITS | NO_HITS | NO_HITS | NO_HITS |
| IGORF_102 | NO_HITS | NO_HITS | NO_HITS | NO_HITS |
| IGORF_103 | NO_HITS | NO_HITS | NO_HITS | NO_HITS |
| IGORF_104 | NO_HITS | NO_HITS | NO_HITS | NO_HITS |
| IGORF_105 | Raphanus sativus U5 spliceosomal RNA LOC130497136 ncRNA | 77% | 0.0007 | 88.64 |
| IGORF_106 | A. thaliana genome | 84% | 3.00E-44 | 91.61 |
| IGORF_107 | NO_HITS | NO_HITS | NO_HITS | NO_HITS |
| IGORF_108 | A. thaliana genome | 96% | 8.00E-55 | 93.46 |
| IGORF_109 | NO_HITS | NO_HITS | NO_HITS | NO_HITS |
| IGORF_110 | A. thaliana genome | 65% | 2.00E-25 | 92.31 |
| IGORF_111 | A. thaliana genome | 97% | 9.00E-66 | 88.65 |
| IGORF_112 | A. thaliana genome | 74% | 2.00E-37 | 87.18 |
| IGORF_113 | NO_HITS | NO_HITS | NO_HITS | NO_HITS |
| IGORF_114 | A lyrata uncharacterized LOC110230800 ncRNA | 100% | 2.00E-11 | 100 |
| IGORF_115 | A. thaliana genome | 100% | 4.00E-12 | 90.62 |
| IGORF_116 | NO_HITS | NO_HITS | NO_HITS | NO_HITS |
| IGORF_117 | NO_HITS | NO_HITS | NO_HITS | NO_HITS |
| IGORF_118 | A. lyrata probable inactive leucine-rich repeat receptor-like protein kinase At3g03770 mRNA | 53% | 3.00E-23 | 100 |
| IGORF_120 | A. thaliana genome | 83% | 4.00E-11 | 92.86 |
| IGORF_122 | NO_HITS | NO_HITS | NO_HITS | NO_HITS |
| IGORF_123 | NO_HITS | NO_HITS | NO_HITS | NO_HITS |

| | | | | |
|---|---|---|---|---|
| IGORF_124 | NO_HITS | NO_HITS | NO_HITS | NO_HITS |
| IGORF_125 | NO_HITS | NO_HITS | NO_HITS | NO_HITS |
| IGORF_126 | A. thaliana miR172c primary transcript gene,<br>Arabidopsis lyrata microRNA aly-MIR172c precursor,<br>Brassica rapa genome | 70%,<br>47%,<br>44% | 2E-34,<br>4.00E-22,<br>4.00E-17 | 96.84,<br>100,<br>96.67 |
| IGORF_129 | NO_HITS | NO_HITS | NO_HITS | NO_HITS |
| IGORF_130 | NO_HITS | NO_HITS | NO_HITS | NO_HITS |
| IGORF_131 | NO_HITS | NO_HITS | NO_HITS | NO_HITS |
| IGORF_132 | A. thaliana genome | 80% | 6.00E-52 | 92.16 |
| IGORF_133 | A. thaliana genome | 98% | 7.00E-38 | 97.96 |
| IGORF_134 | A. thaliana genome,<br>Brassica oleracea HDEM genome | 70%,<br>58% | 4E-26,<br>2.00E-13 | 97.37,<br>92.06 |
| IGORF_138 | NO_HITS | NO_HITS | NO_HITS | NO_HITS |
| IGORF_140 | NO_HITS | NO_HITS | NO_HITS | NO_HITS |
| IGORF_141 | NO_HITS | NO_HITS | NO_HITS | NO_HITS |
| IGORF_143 | NO_HITS | NO_HITS | NO_HITS | NO_HITS |
| IGORF_144 | NO_HITS | NO_HITS | NO_HITS | NO_HITS |
| IGORF_145 | NO_HITS | NO_HITS | NO_HITS | NO_HITS |
| IGORF_148 | NO_HITS | NO_HITS | NO_HITS | NO_HITS |
| IGORF_149 | NO_HITS | NO_HITS | NO_HITS | NO_HITS |
| IGORF_150 | NO_HITS | NO_HITS | NO_HITS | NO_HITS |
| IGORF_151 | NO_HITS | NO_HITS | NO_HITS | NO_HITS |
| IGORF_152 | NO_HITS | NO_HITS | NO_HITS | NO_HITS |
| IGORF_155 | NO_HITS | NO_HITS | NO_HITS | NO_HITS |
| IGORF_156 | NO_HITS | NO_HITS | NO_HITS | NO_HITS |
| IGORF_157 | NO_HITS | NO_HITS | NO_HITS | NO_HITS |
| IGORF_158 | NO_HITS | NO_HITS | NO_HITS | NO_HITS |
| IGORF_159 | NO_HITS | NO_HITS | NO_HITS | NO_HITS |
| IGORF_161 | NO_HITS | NO_HITS | NO_HITS | NO_HITS |
| IGORF_162 | NO_HITS | NO_HITS | NO_HITS | NO_HITS |
| IGORF_163 | NO_HITS | NO_HITS | NO_HITS | NO_HITS |
| IGORF_164 | NO_HITS | NO_HITS | NO_HITS | NO_HITS |
| IGORF_165 | NO_HITS | NO_HITS | NO_HITS | NO_HITS |
| IGORF_167 | A. thaliana genome,<br>Arabidopsis thaliana FASCICLIN-like arabinogalactan protein 8 FLA8 mRNA,<br>A. lyrata fasciclin-like arabinogalactan protein 8 LOC9318081 mRNA | 99%,<br>71%,<br>66% | 2E-33,<br>8.00E-22,<br>5.00E-19 | 98.84,<br>100,<br>100 |
| IGORF_168 | A. thaliana genome | 90% | 1.00E-30 | 91.74 |
| IGORF_169 | NO_HITS | NO_HITS | NO_HITS | NO_HITS |
| IGORF_170 | Camelina sativa uncharacterized LOC109132690 ncRNA | 94% | 0.0003 | 96.88 |
| IGORF_171 | A. thaliana genome | 78% | 5.00E-31 | 93.88 |
| IGORF_172 | NO_HITS | NO_HITS | NO_HITS | NO_HITS |
| IGORF_176 | NO_HITS | NO_HITS | NO_HITS | NO_HITS |
| IGORF_178 | NO_HITS | NO_HITS | NO_HITS | NO_HITS |
| IGORF_177 | NO_HITS | NO_HITS | NO_HITS | NO_HITS |
| IGORF_179 | NO_HITS | NO_HITS | NO_HITS | NO_HITS |
| IGORF_180 | NO_HITS | NO_HITS | NO_HITS | NO_HITS |
| IGORF_181 | A. thaliana genome | 39% | 4.00E-11 | 100 |
| IGORF_183 | NO_HITS | NO_HITS | NO_HITS | NO_HITS |
| IGORF_185 | NO_HITS | NO_HITS | NO_HITS | NO_HITS |
| IGORF_186 | NO_HITS | NO_HITS | NO_HITS | NO_HITS |
| IGORF_189 | NO_HITS | NO_HITS | NO_HITS | NO_HITS |
| IGORF_190 | NO_HITS | NO_HITS | NO_HITS | NO_HITS |
| IGORF_191 | NO_HITS | NO_HITS | NO_HITS | NO_HITS |
| IGORF_192 | NO_HITS | NO_HITS | NO_HITS | NO_HITS |
| IGORF_193 | NO_HITS | NO_HITS | NO_HITS | NO_HITS |
| IGORF_194 | NO_HITS | NO_HITS | NO_HITS | NO_HITS |
| IGORF_195 | NO_HITS | NO_HITS | NO_HITS | NO_HITS |
| IGORF_196 | NO_HITS | NO_HITS | NO_HITS | NO_HITS |
| IGORF_198 | A. lyrata nascent polypeptide-associated complex subunit alpha-like protein 3 LOC9309699 mRNA | 73% | 2.00E-66 | 99.32 |
| IGORF_199 | A. lyrata peroxidase 56 LOC9309780 mRNA | 58% | 1.00E-14 | 100 |
| IGORF_200 | NO_HITS | NO_HITS | NO_HITS | NO_HITS |
| IGORF_201 | NO_HITS | NO_HITS | NO_HITS | NO_HITS |
| IGORF_202 | NO_HITS | NO_HITS | NO_HITS | NO_HITS |
| IGORF_203 | NO_HITS | NO_HITS | NO_HITS | NO_HITS |

| | | | | |
|---|---|---|---|---|
| IGORF_204 | A. thaliana genome,<br>A. thaliana uncharacterized misc_RNA AT5G00435 miscRNA | 52%,<br>52% | 0.000000002,<br>0.000000002 | 91.07,<br>91.07 |
| IGORF_205 | NO_HITS | NO_HITS | NO_HITS | NO_HITS |
| IGORF_206 | NO_HITS | NO_HITS | NO_HITS | NO_HITS |
| IGORF_207 | NO_HITS | NO_HITS | NO_HITS | NO_HITS |
| IGORF_208 | NO_HITS | NO_HITS | NO_HITS | NO_HITS |
| IGORF_210 | A. thaliana genome | 100% | 9.00E-33 | 91.23 |
| IGORF_211 | NO_HITS | NO_HITS | NO_HITS | NO_HITS |
| IGORF_212 | NO_HITS | NO_HITS | NO_HITS | NO_HITS |
| IGORF_214 | NO_HITS | NO_HITS | NO_HITS | NO_HITS |
| IGORF_216 | A. thaliana genome,<br>Arabidopsis thaliana ascorbate peroxidase 3 APX3 mRNA | 92%,<br>75% | 2E-15,<br>2.00E-10 | 95,<br>95.83 |
| IGORF_218 | NO_HITS | NO_HITS | NO_HITS | NO_HITS |
| IGORF_219 | A. thaliana genome | 65% | 3.00E-27 | 97.44 |
| IGORF_222 | A. lyrata uncharacterized LOC9305728 mRNA | 100% | 7.00E-10 | 100 |
| IGORF_223 | NO_HITS | NO_HITS | NO_HITS | NO_HITS |
| IGORF_225 | A. thaliana genome | 100% | 4.00E-48 | 88.57 |
| IGORF_226 | NO_HITS | NO_HITS | NO_HITS | NO_HITS |
| IGORF_227 | NO_HITS | NO_HITS | NO_HITS | NO_HITS |
| IGORF_228 | NO_HITS | NO_HITS | NO_HITS | NO_HITS |
| IGORF_229 | NO_HITS | NO_HITS | NO_HITS | NO_HITS |
| IGORF_230 | NO_HITS | NO_HITS | NO_HITS | NO_HITS |
| IGORF_231 | Arabidopsis thaliana Ribosomal protein L35 AT5G45590 mRNA | 98% | 6.00E-10 | 78.63 |
| IGORF_232 | A. lyrata ankyrin repeat-containing protein ITN1 LOC110225629 mRNA | 100% | 2.00E-27 | 100 |
| IGORF_233 | NO_HITS | NO_HITS | NO_HITS | NO_HITS |
| IGORF_235 | NO_HITS | NO_HITS | NO_HITS | NO_HITS |
| IGORF_236 | NO_HITS | NO_HITS | NO_HITS | NO_HITS |
| IGORF_238 | NO_HITS | NO_HITS | NO_HITS | NO_HITS |
| IGORF_241 | NO_HITS | NO_HITS | NO_HITS | NO_HITS |
| IGORF_242 | NO_HITS | NO_HITS | NO_HITS | NO_HITS |
| IGORF_244 | NO_HITS | NO_HITS | NO_HITS | NO_HITS |
| IGORF_246 | NO_HITS | NO_HITS | NO_HITS | NO_HITS |
| IGORF_247 | NO_HITS | NO_HITS | NO_HITS | NO_HITS |
| IGORF_248 | NO_HITS | NO_HITS | NO_HITS | NO_HITS |
| IGORF_252 | NO_HITS | NO_HITS | NO_HITS | NO_HITS |
| IGORF_251 | NO_HITS | NO_HITS | NO_HITS | NO_HITS |
| IGORF_253 | NO_HITS | NO_HITS | NO_HITS | NO_HITS |
| IGORF_254 | A. lyrata uncharacterized LOC9301140 mRNA,<br>A. thaliana genome | 63%,<br>32% | 6E-46,<br>1.00E-12 | 100,<br>96.15 |

**Table S4:** Supplementary Megablast results for each translated *de novo* ORF (core nucleotide database nt, update date: 2025/07/31), with default search parameters

| Species | Population | GPS coordinates | Sample accession | Reference |
|---|---|---|---|---|
| *A. halleri* | Harz Moutains (Germany) | 51.6655, 11.1584 | SAMEA115474513 | *Suda et al., 2024* |
| *A. halleri* | Western Carpathians (Slovakia) | 48.8564, 20.9203 | SAMEA115474514 | *Suda et al., 2024* |
| *A. halleri* | Western Carpathians (Slovakia) | 48.7658, 21.1297 | SAMEA115474515 | *Suda et al., 2024* |
| *A. halleri* | Harz Moutains (Germany) | 51.8626, 10.3001 | SAMEA115474516 | *Suda et al., 2024* |
| *A. halleri* | Siegerland (Germany) | 51.0054, 8.0066 | SAMEA115474517 | *Suda et al., 2024* |
| *A. halleri* | Kosice-okolie District (Slovakia) | 48.7523, 20.8964 | SAMEA115474518 | *Suda et al., 2024* |
| *A. halleri* | Miasteczko Śląskie (Poland) | 50.5035, 18.9357 | SAMEA115474519 | *Suda et al., 2024* |
| *A. halleri* | Miasteczko Śląskie (Poland) | 50.5035, 18.9357 | SAMEA115474520 | *Suda et al., 2024* |
| *A. halleri* | Bayern (Germany) | 50.4114, 11.5553 | SAMEA115474521 | *Suda et al., 2024* |
| *A. halleri* | Sauerland (Germany) | 51.3064, 8.4852 | SAMEA115474522 | *Suda et al., 2024* |
| *A. halleri* | Tatra Mountains (Poland) | 49.2792, 19.9661 | SAMEA115474523 | *Suda et al., 2024* |
| *A. halleri* | Tatra Mountains (Poland) | 49.2792, 19.9661 | SAMEA115474524 | *Suda et al., 2024* |
| *A. halleri* | Maramures County (Romania) | 47.8085, 23.6171 | SAMEA115474525 | *Suda et al., 2024* |
| *A. halleri* | Suceava County (Romania) | 47.5277, 25.4276 | SAMEA115474526 | *Suda et al., 2024* |
| *A. halleri* | Maramures County (Romania) | 47.6947, 23.7769 | SAMEA115474527 | *Suda et al., 2024* |
| *A. halleri* | Cluj County (Romania) | 46.8632, 22.8077 | SAMEA115474528 | *Suda et al., 2024* |
| *A. halleri* | Lana di Dentro (Italy) | 46.4931, 10.8895 | SAMN43174266 | *Pavan et al. 2025* |
| *A. halleri* | Ponte Nossa (Italy) | 45.8593, 9.8767 | SAMN42903595 | *Pavan et al. 2025* |
| *A. halleri* | Paisco Loveno (Italy) | 46.0556, 10.2429 | SAMN42903597 | *Pavan et al. 2025* |
| *A. halleri* | Mortagne-du-Nord (France) | 50.4938, 3.4563 | SAMN43174278 | *Pavan et al. 2025* |
| *A. halleri* | Wallenfels (Germany) | 50.4114, 11.5553 | SAMN42903598 | *Pavan et al. 2025* |
| *A. halleri* | Wallenfels (Germany) | 50.4114, 11.5553 | SAMN42903599 | *Pavan et al. 2025* |

| | | | | |
|---|---|---|---|---|
| *A. halleri* | Falkenstein (Germany) | 51.6656, 11.1585 | SAMN43174267 | *Pavan et al. 2025* |
| *A. halleri* | Langelsheim (Germany) | 51.9488, 10.3488 | SAMN42903593 | *Pavan et al. 2025* |
| *A. halleri* | Croglio (Switzerland) | 45.9875, 8.8395 | SAMN43174284 | *Pavan et al. 2025* |
| *A. halleri* | Zocca-Vicosoprano (Switzerland) | 46.3716, 9.6601 | SAMN43174279 | *Pavan et al. 2025* |
| *A. halleri* | Kowari (Poland) | 50.7586, 15.8467 | SAMN42903592 | *Pavan et al. 2025* |
| *A. halleri* | Zakopane (Poland) | 49.2792, 19.9660 | SAMN42914077 | *Pavan et al. 2025* |
| *A. halleri* | Miasteczko Śląskie (Poland) | 50.5035, 18.9357 | SAMN42903594 | *Pavan et al. 2025* |
| *A. halleri* | Blidari (Romania) | 47.8085, 23.6171 | SAMN43174271 | *Pavan et al. 2025* |
| *A. halleri* | Munții Gutâi (Romania) | 47.6947, 23.7769 | SAMN43174272 | *Pavan et al. 2025* |
| *A. halleri* | Tranisu (Romania) | 46.8632, 22.8077 | SAMN43174277 | *Pavan et al. 2025* |
| *A. halleri* | Fundu Moldovei (Romania) | 47.5277, 25.4276 | SAMN43174273 | *Pavan et al. 2025* |
| *A. halleri* | Galmeikogel - Annarotte (Austria) | 47.8485, 15.3902 | SAMN43174275 | *Pavan et al. 2025* |
| *A. halleri* | Bad Gastein (Austria) | 47.1340, 13.1267 | SAMN43174276 | *Pavan et al. 2025* |
| *A. halleri* | Castle Rabenstein (Austria) | 47.2496, 15.3089 | SAMN43174268 | *Pavan et al. 2025* |
| *A. halleri* | Steinbachrotte (Austria) | 47.9383, 15.4339 | SAMN43174287 | *Pavan et al. 2025* |
| *A. lyrata* | Pinery Provincial Park (North America) | 43.2640, -81.8397 | SAMN43781242 | *Pavan et al. 2025* |
| *A. lyrata* | Pinery Provincial Park (North America) | 43.2640, -81.8397 | SAMN43781218 | *Pavan et al. 2025* |
| *A. lyrata* | Pinery Provincial Park (North America) | 43.2640, -81.8397 | SAMN43781219 | *Pavan et al. 2025* |
| *A. lyrata* | Rondeau Provincial Park (North America) | 42.2613, -81.8464 | SAMN43781240 | *Pavan et al. 2025* |
| *A. lyrata* | Rondeau Provincial Park (North America) | 42.2614, -81.8464 | SAMN43781237 | *Pavan et al. 2025* |
| *A. lyrata* | Rondeau Provincial Park (North America) | 42.2614, -81.8464 | SAMN43781234 | *Pavan et al. 2025* |
| *A. lyrata* | Tobermory georgian bay (North America) | 45.2417, -81.5175 | SAMN43781213 | *Pavan et al. 2025* |
| *A. lyrata* | Tobermory georgian bay (North America) | 45.2417, -81.5175 | SAMN43781216 | *Pavan et al. 2025* |

| | | | | |
|---|---|---|---|---|
| *A. lyrata* | Tobermory georgian bay (North America) | 45.2417, -81.5175 | SAMN43781214 | *Pavan et al. 2025* |
| *A. lyrata* | Tobermory singing sands (North America) | 45.1925, -81.5839 | SAMN43781247 | *Pavan et al. 2025* |
| *A. lyrata* | Tobermory singing sands (North America) | 45.1925, -81.5839 | SAMN43781227 | *Pavan et al. 2025* |
| *A. lyrata* | Tobermory singing sands (North America) | 45.1925, -81.5839 | SAMN43781228 | *Pavan et al. 2025* |
| *A. lyrata* | ndiana Dune National Lakeshore (North America) | 41.6689, -87.0422 | SAMN43781217 | *Pavan et al. 2025* |
| *A. lyrata* | ndiana Dune National Lakeshore (North America) | 41.66897, -87.04221 | SAMN43781221 | *Pavan et al. 2025* |
| *A. lyrata* | ndiana Dune National Lakeshore (North America) | 41.6689, -87.0422 | SAMN43781222 | *Pavan et al. 2025* |
| *A. lyrata* | Storr (UK) | 57.50, -6.16 | SAMN06141186 | *Mattila et al. 2017* |
| *A. lyrata* | Storr (UK) | 57.50, -6.16 | SAMN06141185 | *Mattila et al. 2017* |
| *A. lyrata* | Stubbsand (Sweden) | 63.22, 18.95 | SAMN06141184 | *Mattila et al. 2017* |
| *A. lyrata* | Stubbsand (Sweden) | 63.22, 18.95 | SAMN06141183 | *Mattila et al. 2017* |
| *A. lyrata* | Stubbsand (Sweden) | 63.22, 18.95 | SAMN06141182 | *Mattila et al. 2017* |
| *A. lyrata* | Stubbsand (Sweden) | 63.22, 18.95 | SAMN06141181 | *Mattila et al. 2017* |
| *A. lyrata* | Plech (Germany) | 49.63, 11.51 | SAMN06141187 | *Mattila et al. 2017* |
| *A. lyrata* | Plech (Germany) | 49.63, 11.51 | SAMN06141188 | *Mattila et al. 2017* |
| *A. lyrata* | Plech (Germany) | 49.63, 11.51 | SAMN06141189 | *Mattila et al. 2017* |
| *A. lyrata* | Mayodan (USA) | 36.42, -79.97 | SAMN06141193 | *Mattila et al. 2017* |
| *A. lyrata* | Mayodan (USA) | 36.42, -79.97 | SAMN06141194 | *Mattila et al. 2017* |
| *A. lyrata* | Mayodan (USA) | 36.42, -79.97 | SAMN06141195 | *Mattila et al. 2017* |
| *A. lyrata* | Spiterstulen (Norway) | 61.63, 8.4 | SAMN06141173 | *Mattila et al. 2017* |
| *A. lyrata* | Spiterstulen (Norway) | 61.63, 8.4 | SAMN06141174 | *Mattila et al. 2017* |
| *A. lyrata* | Spiterstulen (Norway) | 61.63, 8.4 | SAMN06141175 | *Mattila et al. 2017* |
| *A. lyrata* | Austria | 47.9015, 15.9686 | SAMEA5947868 | *Marburger et al. 2019* |

| | | | | |
|---|---|---|---|---|
| *A. lyrata* | Austria | 47.9015, 15.9686 | SAMEA5947869 | *Marburger et al. 2019* |
| *A. lyrata* | Austria | 47.92251, 15.98176 | SAMEA5947870 | *Marburger et al. 2019* |
| *A. lyrata* | Austria | 47.92251, 15.98176 | SAMEA5947871 | *Marburger et al. 2019* |
| *A. lyrata* | Plech (Germany) | 49.65, 11.45 | SAMEA6943357 | *Takou et al. 2021* |
| *A. lyrata* | Plech (Germany) | 49.65, 11.45 | SAMEA6943358 | *Takou et al. 2021* |
| *A. lyrata* | Plech (Germany) | 49.65, 11.45 | SAMEA6943359 | *Takou et al. 2021* |
| *A. lyrata* | Spiterstulen (Norway) | 61.62, 8.40 | SAMN09069416 | *Hämälä et al. 2018* |
| *A. lyrata* | Spiterstulen (Norway) | 61.62, 8.40 | SAMN09069417 | *Hämälä et al. 2018* |
| *A. lyrata* | Spiterstulen (Norway) | 61.62, 8.40 | SAMN09069418 | *Hämälä et al. 2018* |

**Table S5:** Summary of the population whole-genome sequencing data
